# Long-read sequencing maps transposable element variation and its regulatory and epigenetic effects in the human brain

**DOI:** 10.64898/2026.07.02.735893

**Authors:** Alexis Ayuketah, Melissa Meredith, Cristian Groza, Jon Moller, Kensuke Daida, Adam Catching, Cory Weller, Cedric Kouam, Luis Paulin, Laksh Malik, Breeana Baker, Fangle Hu, Sarah Bromberek, Pilar Alvarez Jerez, Kimberly Paquette, Michal Izydorczyk, Bida Gu, Mark J. P. Chaisson, Ben Middlehurst, Vivien J. Bubb, John P. Quinn, Emma Price, Andrew B. Singleton, Miten Jain, Cornelis Blauwendraat, Mike Nalls, Mark R Cookson, Xylena Reed, Fritz J. Sedlazeck, Clément Goubert, Kimberley J. Billingsley

**Affiliations:** Center for Alzheimer’s and Related Dementias, National Institute on Aging and National Institute of Neurological Disorders and Stroke, National Institutes of Health, Bethesda, MD, USA; DataTecnica LLC, Washington, DC, USA; Montreal Heart Institute, Université de Montréal, QC, Canada; Laboratory of Neurogenetics, National Institute on Aging, National Institutes of Health, Bethesda, MD, USA; Human Genome Sequencing Center, Baylor College of Medicine, Houston, TX, USA; Department of Neurodegenerative Disease, UCL Queen Square Institute of Neurology, University College London, London, UK; Department of Biology, Johns Hopkins University, Baltimore, MD, USA; Global Parkinson’s Genetics Program (GP2), Chevy Chase, MD, USA; Department of Quantitative and Computational Biology, University of Southern California, Los Angeles, CA, USA; Department of Pharmacology and Therapeutics, University of Liverpool, Liverpool, UK; Molecular Pathology Section, Laboratory of Immunogenetics, National Institute of Allergy and Infectious Diseases, NIH; Department of Bioengineering, Department of Physics, Northeastern University, Boston, MA, USA; Center for Cancer Research, National Cancer Institute, National Institutes of Health, USA; R. Ken Coit College of Pharmacy, University of Arizona, Tucson, AZ, USA

## Abstract

Transposable elements (TEs) are mobile DNA sequences that shape genome architecture and gene regulation, yet their roles in the human brain remain largely unresolved. Short-read sequencing lacks the resolution to accurately map TE insertions, detect associated structural variants, and resolve highly repetitive regions. Here, we leverage long-read whole-genome sequencing to profile germline TE insertions in postmortem brain tissue from two ancestrally diverse cohorts: the North American Brain Expression Consortium (NABEC; European ancestry, *n* = 205) and the Human Brain Collection Core (HBCC; African and African-admixed ancestry, *n* = 146). We identified 2,842 and 1,660 high-confidence non-reference insertions in HBCC and NABEC, respectively, spanning *Alu*, LINE-1, and SVA elements. We then also further characterized complex short tandem repeat and variable number tandem repeat variation within reference SVA and *Alu* loci. Reference TEs were also found to mediate complex structural variants at loci implicated in brain development and neurodegenerative disease, with several showing ancestry-specific patterns. Integration of bulk RNA-sequencing data identified TE expression quantitative trait loci, including insertions that modulate neuronal gene expression. Single-nucleus RNA sequencing revealed cell-type-specific effects of TE regulation across cortical populations. Long-read methylation profiling further demonstrated age-associated epigenetic regulation of both reference and non-reference *Alu* elements. As a community resource, we release a catalog of TE insertions, allele frequencies, and ancestry-specific distributions to enable future functional and disease-focused investigations. Together, these findings highlight the widespread regulatory and epigenetic influence of TEs in the human brain and establish long-read sequencing as a powerful approach for uncovering cell-type- and population-specific TE dynamics.

## Introduction

Discovered by Barbara McClintock in maize in the 1940s (*1*), transposable elements (TEs) are DNA sequences that can “move” within the genome. TEs are classified into two major groups: retrotransposons and DNA transposons. Retrotransposons represent the primary active elements in the human genome and mobilize via an RNA intermediate (*2*). This category of TEs includes short interspersed nuclear elements (SINEs), such as *Alu*, SINE-VNTR-*Alu* (SVA), long interspersed nuclear element-1 (LINE-1 or L1), and human endogenous retroviruses (HERVs) (*3*), in which these repetitive sequences comprise two-thirds of the human genome (4). SINEs and SVAs are non-autonomous, requiring another TE to be able to move, while LINE-1s are autonomous, providing all the machinery needed for movement (via target-primed reverse transcription (TPRT)). These unique mechanisms of mobilization underpin the emergence of novel TE insertions, contributing significantly to genetic diversity.

TE variation can be broadly categorized into reference and non-reference elements. Reference TEs are elements already annotated in the GRCh38 human reference genome by RepeatMasker. Variation in these elements can arise either through canonical deletions, leading to presence/absence polymorphisms, or through alterations in their internal repeat domains, such as changes in the length of variable number tandem repeats (VNTRs). Non-reference TEs also represent presence/absence variation, where insertions are present in individual genomes but absent from the GRCh38 reference genome. While the overall frequency of polymorphic germline TE non-reference insertions is relatively low in humans, their distribution varies across ancestral populations, with approximately 25% of polymorphic TE loci being population-specific (5). In line with generally observed diversity patterns, genome-wide analyses of polymorphic TEs show that African populations harbour the highest levels of TE genetic diversity, with lower diversity in non-African populations (*4*). Among non-African groups, relative diversity patterns vary by TE family, with polymorphic TE-wide analyses showing lower diversity in Asian and European populations (*4*), whereas *Alu*-specific studies report the lowest *Alu* diversity in European populations (*5*).

TE integration into the genome can influence gene expression and genome stability (*6*, *7*). Epigenetically, they are sites of significant CpG methylation (*8*) and can form G-quadruplexes due to their guanine-rich nature (*9*). Transcriptionally, they can both act as and integrate into cis-regulatory elements (*10*). Post-transcriptionally, they can be incorporated as cryptic exons (*11*), alter gene products, or be expressed as and/or regulate ncRNAs (*12*). In some cases, these insertions have been directly implicated in disease. For instance, X-linked dystonia-parkinsonism is associated with an SVA insertion in *TAF1* within Filipino male populations (*13*). TE-mediated disruptions have also been reported in amyotrophic lateral sclerosis (ALS) (*14–16*), Alzheimer’s disease (AD) (*17*, *18*), and Parkinson’s disease (PD) (*19–21*). However, the precise mechanisms through which TEs contribute to these neurodegenerative diseases remain poorly understood.

Despite their significance, TE detection and accurate genotyping remain challenging due to the highly repetitive nature of these elements. Short-read sequencing technologies often fail to resolve these elements, leading to misplacement, incomplete annotation, and low sensitivity (*22*), resulting in less precise genotyping information, which is crucial for association studies. The advent of long-read sequencing technologies, including Pacific Biosciences and Oxford Nanopore Technologies (ONT), offers a promising solution. Due to their significantly longer read lengths (multiple kilobases), these platforms enable more accurate breakpoint resolution, facilitate de novo assembly, and improve TE annotation, even at lower genome coverage (*22*). However, despite these advances, tools for robust TE annotation and characterization remain limited. Many current computational tools, such as MELT (*23*), Tangram (*24*), and TEMP (*24*, *25*), have been developed to analyze polymorphic TEs in short-read datasets (23). However, there are discrepancies in their outputs, underscoring ongoing limitations in TE detection and furthering the need for scalable methodologies that leverage the resolution of long-read sequencing.

The most direct attempt to profile TEs in the human brain using long-read sequencing comes from Ramirez et al. (*26*), who analyzed 18 postmortem frontal cortex samples at an average coverage of 7.4×, identifying over 1,000 novel non-reference TEs, predominantly *Alu*, followed by LINE-1, SVA, and HERV-K. These elements were enriched in intronic and regulatory regions and exhibited differential DNA methylation patterns, with younger *Alu* and LINE-1 insertions showing higher CpG methylation levels than older elements. However, due to the small sample size and limited integration with functional genomic data, the broader impact of polymorphic TEs on gene regulation and brain biology remains largely unexplored.

Here, we present the most comprehensive survey to date of TE diversity in the human genome using long-read sequencing. We leverage ONT long-read whole-genome sequencing data from 351 individuals of European and African ancestries, including samples from the North American Brain Expression Consortium (NABEC) and the Human Brain Collection Core (HBCC). From here, we systematically identify and annotate polymorphic insertions from three active TE families, *Alu*, SVA, and LINE-1. We explore the role of TEs in shaping structural variation, including complex and rare genomic rearrangements; evaluate their regulatory effects on gene expression through quantitative trait locus (QTL) analyses; and investigate DNA methylation landscapes at TE loci to uncover patterns of epigenetic regulation. Together, these analyses provide a comprehensive population-scale reference of TE variation and function in the human brain, establishing a foundational resource for future studies of TE biology and their role in health and disease.

## Results

### TEs are abundant and polymorphic across individuals and populations

To systematically assess TE presence/absence, we used previously generated long-read sequencing data from Billingsley et al. (*27*) for two post-mortem brain cohorts (NABEC and HBCC, n = 351 total). Reads had already been basecalled, aligned to GRCh38, and provided as mapped BAM files (*27*). These were then input into GraffiTE (*28*), a Nextflow-based pipeline for detecting, annotating, and genotyping polymorphic TEs from long-read sequencing data. GraffiTE integrates high-performing tools, including Sniffles2, RepeatMasker, and GraphAligner, within a containerized environment, enabling reproducible and scalable analyses across multiple samples. Notably, GraffiTE can detect and report both TE insertions (non-reference) and deletions (reference TE), a feature not commonly implemented in comparable methods (*28*). For an overview of the TE genotype generation in this present study and downstream analyses, see **Figure 1**.

**Figure 1.**
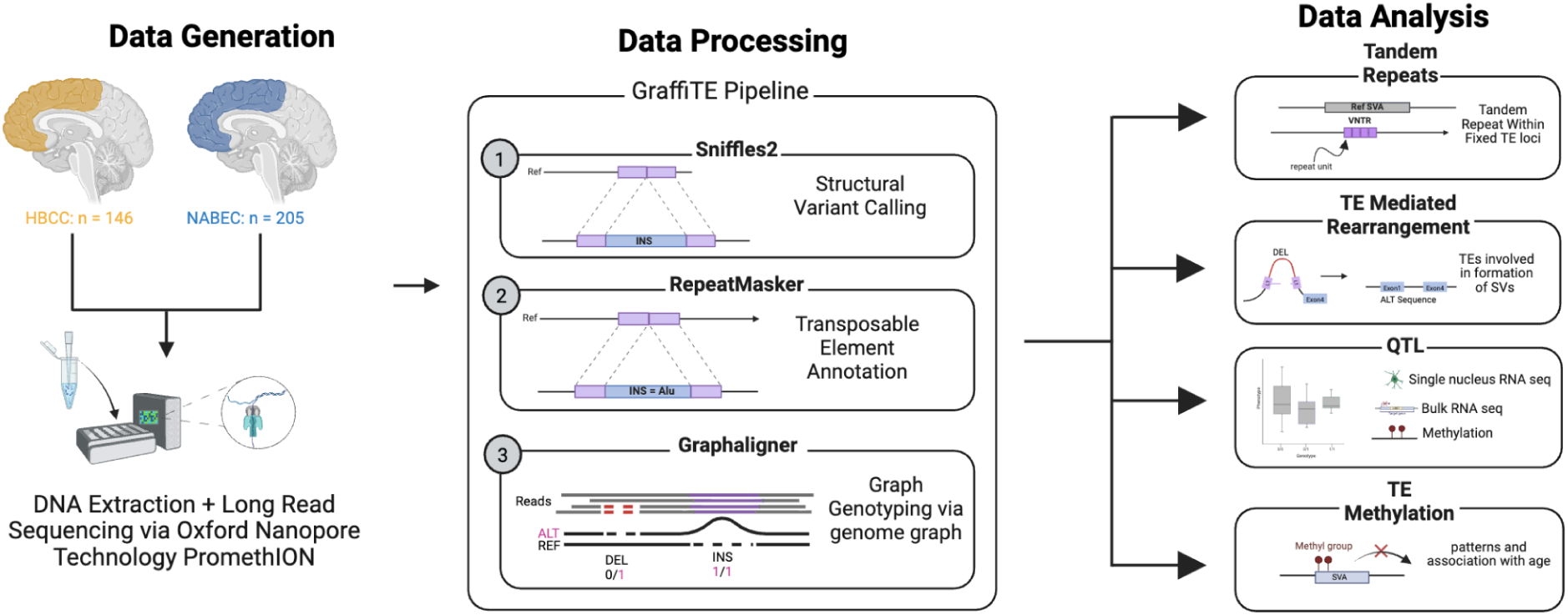
Overview of TE characterization and downstream analyses. *Left*: DNA was extracted from postmortem frontal cortex samples from two cohorts: NABEC (European) and HBCC (African + African Admixed), and sequenced using ONT. *Middle*: Reads mapped to GRCh38 were processed with the GraffiTE pipeline for TE annotation and genome graph-based genotyping. *Right*: Resulting TE variant calls were integrated into downstream analyses: (1) tandem repeat analysis to further characterize TE repeat length variability via VAMOS, (2) detection of TE-mediated SV formation, (3) QTL mapping to assess TE effects on gene expression and DNA methylation, and (4) assessment of TE methylation landscapes, including age-associated changes.

Across both the NABEC and HBCC cohorts, GraffiTE detected 19,699 unique TE candidates. Here, unique refers to a TE variant that is found at least once across the cohorts. Of these, the majority were reference TEs (12,409 (62.9%)), while the rest were non-reference candidate TEs (7,290 (37.1%)). Of the reference TEs, 57.6% were population-specific in HBCC, while 43.5% were population-specific in NABEC. For non-reference, 69.7% and 58.1% were population-specific in HBCC and NABEC, respectively.

To generate a set of high-confidence TEs with the presence/absence genotypes and to maintain consistency with previous reports, we applied stringent filtering to annotate one TE per structural variant. We retained structural variants larger than 250 bp with a single TE annotation spanning at least 80% of the sequence (two annotations were retained if marked as a LINE-1 5′ inversion, as recommended in the GraffiTE documentation). Of these, we only retained TEs belonging to three classes: *Alu*, LINE-1, and SVA.

Most of the high-confidence set of reference TE variants were common (allele frequencies (AF) higher than 5%) (HBCC=67.2%, NABEC=71.5%). Across both cohorts, we identified 3,624 unique TEs, comprising 2,151 and 1,797 *Alu* elements, 803 and 530 LINE-1 elements, and 74 and 68 SVA elements, in HBCC and NABEC, respectively. Interestingly, most of these reference TE variants were intronic: ∼65% of *Alu*s, 58% of LINE-1s, and 67% of SVAs, aligning with prior TE studies (*26*). Per genome, in the HBCC, we found an average of 1096 *Alu*, 213 LINE-1, and 42 SVA. In the NABEC, we identified 947 *Alu*, 182 LINE-1, and 33 SVA per genome (Fig. 2, Supplementary Table 2).

**Figure 2.**
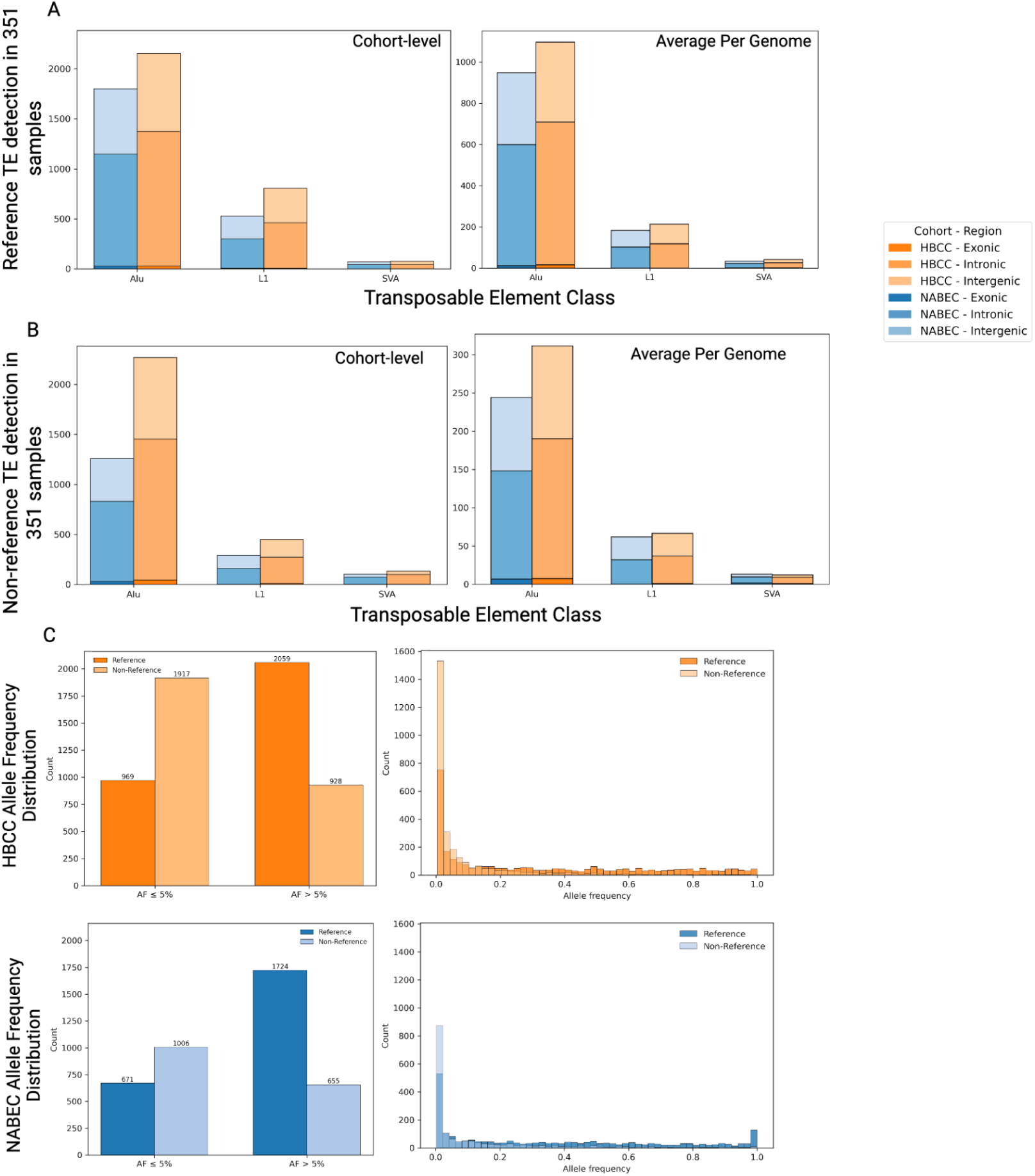
Filtered counts of reference and non-reference across two different populations. (A) Reference TE characterization, broken down by genic region and TE subtype, for both total TEs found (*left*) and average counts per genome for both HBCC and NABEC (*right*). (B) Non-reference TE characterization, broken down by genic region and TE subtype, for both total TEs found (*left*) and average counts per genome for both HBCC and NABEC (*right*). (C) Allele Frequency of TEs detected by reference and non-reference for HBCC (*top*) and NABEC (*bottom*). *Left*: Allele Frequency distribution. *Right*: Allele Frequency counts by rare (≤ 5%) and common (> 5%).

As expected, the majority of the high confidence non-reference TEs were rare (AF ≤5%; HBCC= 59.5% and NABEC =54.6%). Across both cohorts, we found 3,675 unique TEs, comprising 2,264 and 1,259 *Alu*, 447 and 295 LINE-1, and 134 and 107 SVA, in HBCC and NABEC, respectively. Similar to reference TEs, most non-reference elements were also located in intronic regions, including 65.1% of *Alu*, 59.9% of LINE-1, and 73.3% of SVA insertions. Per genome, we identified an average of 311 and 243 *Alu*, 65 and 61 LINE-1, and 12 and 13 SVA, in HBCC and NABEC, respectively. These counts align with prior population-scale long-read studies, which report similar numbers of polymorphic insertions per genome (25). Across both reference and non-reference categories, *Alu* elements were the most abundant TE type, and the HBCC cohort consistently harbored more TE variants than NABEC (Fig. 2A–B, Supplementary Table 2), indicating that TEs are polymorphic and contain ancestry-specific genetic variation.

### Reference SVAs exhibit broad genomic distribution and complex allelic variation

While the above analysis captures presence/absence polymorphism relative to the reference genome, it does not resolve variation within reference TEs, particularly within the tandem-repeat domains of SVA elements. Thus, to complement, we separately analyzed the variation within the VNTR and STR domains of reference SVA elements. Using VAMOS, a tandem repeat annotation tool, we systematically characterized variability in repeat length across cohorts. This tool allows for the identification of the precise number of repeat units (length) and the specific motif composition (sequence) of the repeat, going beyond simple presence/absence analysis.

Repeat length variability is a near-universal feature of reference SVA elements in the human genome. Of the 5,398 reference SVA elements in GRCh38, approximately 74% harbored polymorphic STR domains and ∼62% harbored polymorphic VNTR domains across individuals in both cohorts, demonstrating that the majority of reference SVAs are not fixed in repeat composition but instead constitute highly dynamic loci. Both domain types were extensively multiallelic. STR loci reached up to 380 and 292 alternate alleles in NABEC and HBCC, respectively, with cohort-wide means of 74.66 ± 99.16 and 42.45 ± 55.99 alleles per locus, while VNTR loci were even more complex, with ∼99% being multiallelic with mean counts of 56.60 ± 71.44 and 34.06 ± 40.98 in NABEC and HBCC, respectively. The elevated allele counts in NABEC relative to HBCC most likely reflect a technical rather than biological difference: R9.4.1 sequencing chemistry, used for NABEC, carries higher raw error rates than the R10.4.1 platform used for HBCC, and because VAMOS does not collapse closely related repeat length genotypes, allelic inflation is expected in R9-derived data.

At the level of total repeat length, the two cohorts were strikingly concordant. STR domains spanned 2–15,610 bp in HBCC and 2–15,687 bp in NABEC, with near-identical means of 85.52 ± 212.63 bp and 84.75 ± 212.61 bp. VNTR domains were substantially more expanded, with mean lengths exceeding 1.1 kb in both cohorts (HBCC: 1,133.76 ± 1,013.70 bp; NABEC: 1,138.50 ± 1,032.04 bp) and maximum lengths of 13.7 kb and 18.4 kb, respectively. This range of structural variation, spanning orders of magnitude within a single TE family, is consistent with SVAs representing one of the most structurally diverse classes of repetitive elements in the human genome, as previously evidenced by the extraordinarily high rate of internal structural variability documented at SVA loci relative to all other TE classes (*29*).

Having resolved per-sample SVA repeat lengths using VAMOS, we next asked whether this variation mapped to genomic loci with established links to neurodegeneration risk, and whether it had detectable effects on local gene expression. Variable reference SVA elements were identified across multiple AD and PD risk loci in both cohorts, including *ACE, APOE,* and *BIN1*, suggesting that repeat-length variation at these sites contributes to regulatory signals at disease-associated loci that are otherwise invisible to standard variant-calling approaches. To test this directly, we performed linear regression of total repeat length against RNA expression, identifying a subset of significant associations at AD/PD loci. VNTR domains yielded two and four significant expression associations in HBCC and NABEC, respectively, implicating *KANSL1, HLA-DRB5,* and *HLA-DRB6*, while STR domains yielded one NABEC association at *KANSL1-AS1*. The most compelling example was a reference SVA VNTR located approximately 55 kb and 59 kb upstream of *KANSL1* and *KANSL1-AS1*, respectively, where greater repeat length was robustly associated with increased expression of both genes (*KANSL1*: p = 3.44×10^−6^; *KANSL1-AS1*: p = 9.72×10^−28^; Fig. 3).

**Figure 3.**
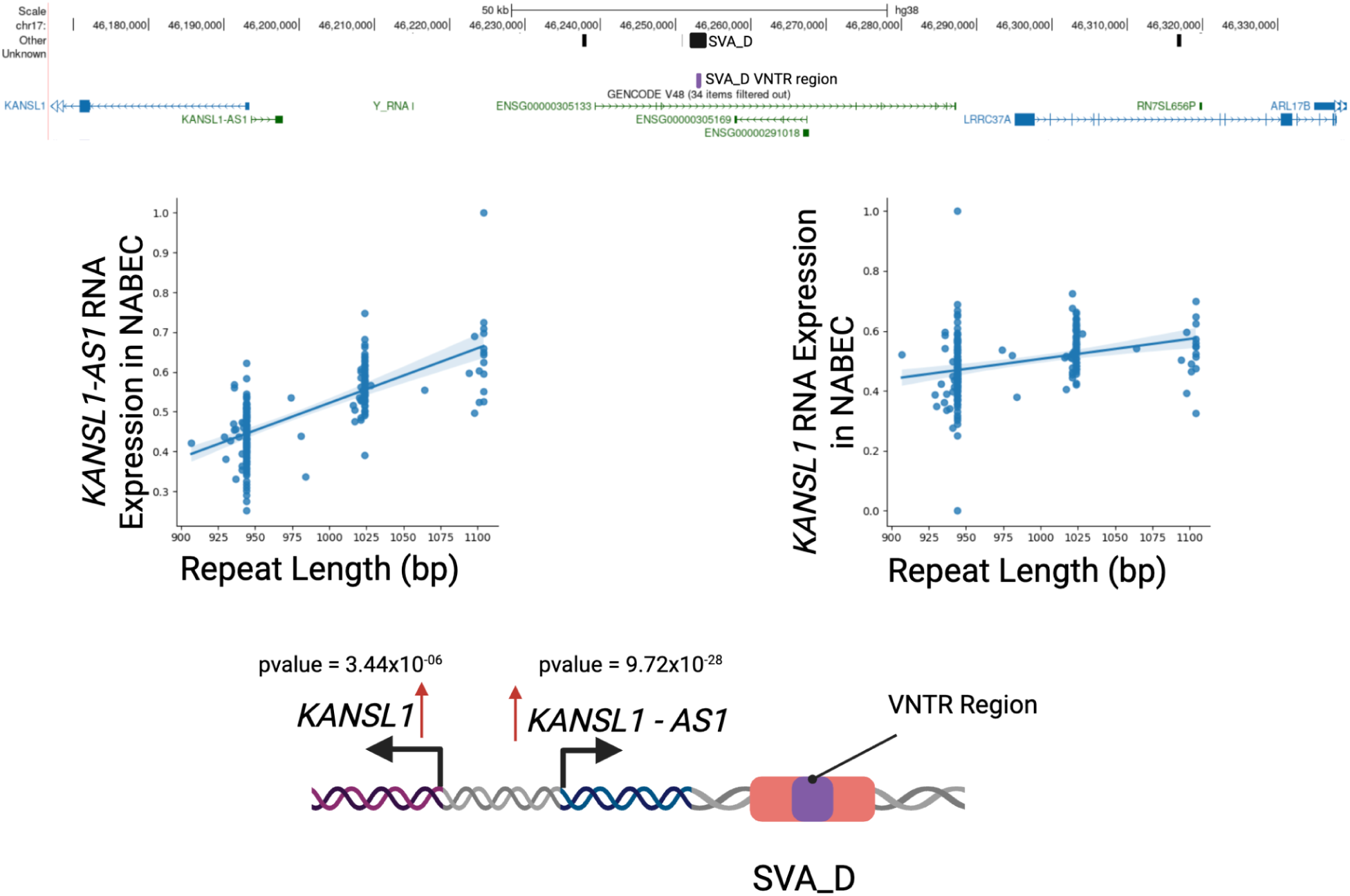
VNTR domain within a reference SVA element ∼50-60 Kb upstream from *KANSL1* and *KANSL1-AS1* locus is associated with higher expression in NABEC. Top: UCSC Genome Browser Image of VNTR region. Middle: Regression line of total repeat length and RNA expression in both *KANSL1* and *KANSL1-AS1*. Bottom: Cartoon schematic of the impact of the VNTR region repeat length variability on gene expression.

Collectively, these findings demonstrate that reference SVA elements harbor extensive repeat-length variability that may be functionally relevant, with repeat length at select loci significantly associated with gene expression in AD and PD risk genes. This suggests that variability in repeat length may be associated with regulatory function in the human genome and warrants deeper investigation.

### Complex allelic patterns and extensive genomic distribution of reference *Alu* variation

*Alu* elements are composed of two monomers separated by a polymorphic middle A-rich region and a poly-A tail of variable length, sequences that act as a cradle for STR formation. Mutations disrupting the poly-A tail can generate STRs that are prone to expansion, with pathogenic alleles capable of producing toxic repeat RNAs or interfering with local gene expression (*30*, *31*). We therefore extended our VAMOS analysis to characterise STR and VNTR variability within reference *Alu* elements. Across more than one million reference *Alu* elements in GRCh38, VAMOS identified repeat-length variability in 82.3% of STR domains and 7.7% of VNTR domains across both cohorts. The majority of variable domains were multiallelic, with maximum alternate allele counts of 292 in HBCC and 380 in NABEC for both repeat types. Mean allele counts were higher in NABEC than HBCC for both STRs (20.56 ± 42.06 vs 14.03 ± 25.02) and VNTRs (23.55 ± 38.82 vs 13.68 ± 21.20). Total repeat length varied widely across *Alu* elements. STR domains had mean lengths of 35.99 ± 106.93 bp in HBCC and 36.20 ± 104.30 bp in NABEC, while VNTR domains were substantially more expanded, with mean lengths of 393.24 ± 540.59 bp in both cohorts. The greater apparent variability in NABEC relative to HBCC, consistent across both repeat classes and their length distributions, likely reflects the higher per-read error rate around repetitive sequences associated with R9 chemistry in NABEC, as observed for SVA elements, rather than true biological differences between cohorts.

We next intersected *Alu* elements harbouring variable STR or VNTR sequences with loci previously implicated in AD and PD, identifying 2,244 variable-length STRs and 1,524 variable-length VNTRs overlapping risk loci across both cohorts, including regions near *MAPT-AS1, KANSL1, and APOE*. To evaluate potential regulatory effects, we tested associations between repeat length and local gene expression. VNTR domains yielded a limited number of significant associations at AD/PD loci, one in HBCC and seventeen in NABEC, while STR domains produced a larger number, with one in HBCC and fifty-four in NABEC. The most compelling example was a reference *Alu* STR in intron 2 of *KANSL1*, where greater repeat length was significantly associated with increased gene expression (p = 4.39×10^−14^; Fig. 4).

**Figure 4.**
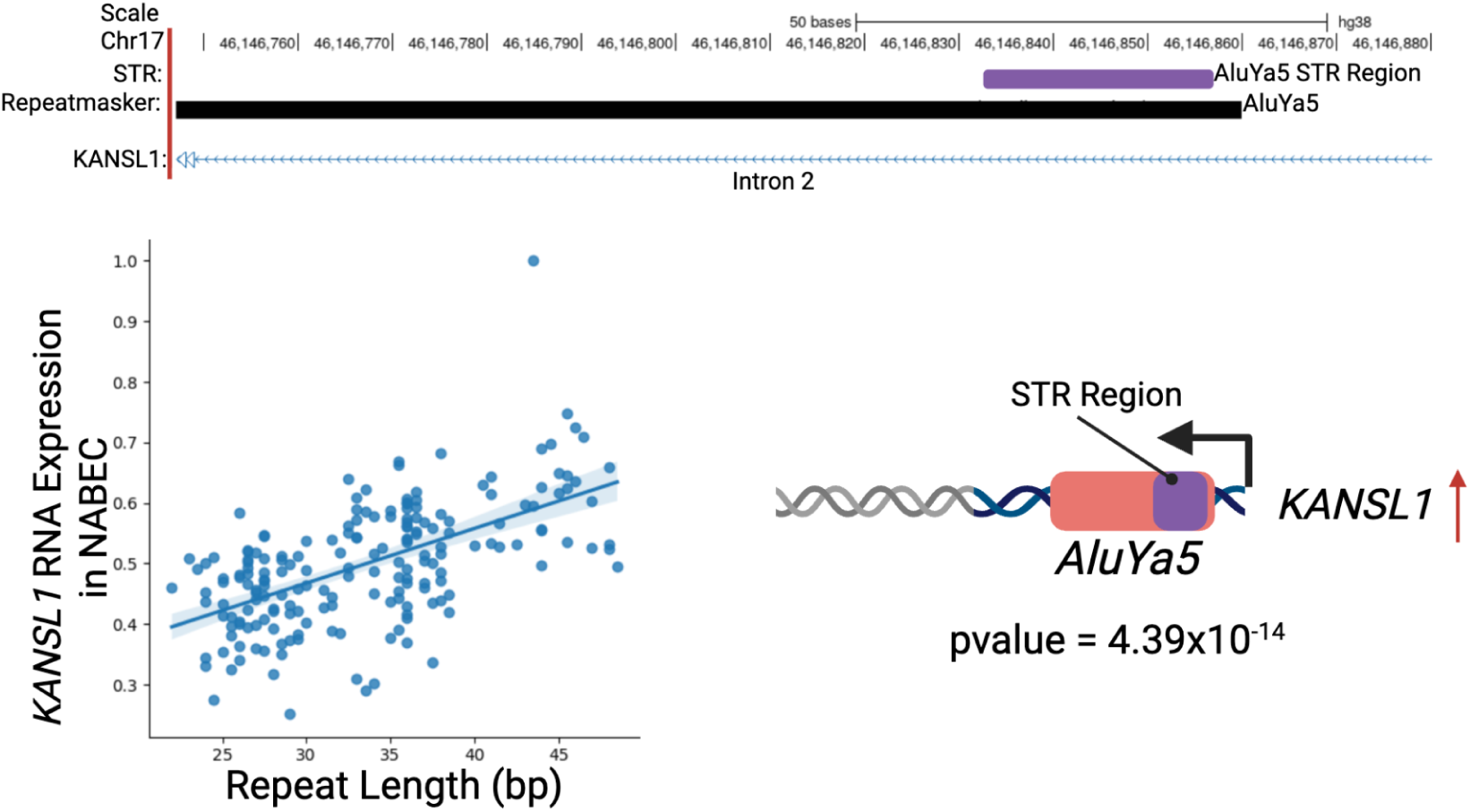
Length of an Intronic STR within a reference *Alu* element at the *KANSL1* locus is associated with increased expression in NABEC. Top: UCSC Genome Browser Image of STR region within intron 2 of *KANSL1*. Bottom Left: Regression Line of total repeat length and RNA expression in *KANSL1*. Bottom Right: Cartoon Schematic of the impact of STR region repeat length variability on gene expression.

### Transposable elements drive complex rate structural variant events across the human genome

We assessed the contribution of TEs to structural variants by identifying TE–mediated rearrangement (TEMR) events. Following the definition by Balachandran et al. (*32*), TEMRs arise when two homologous TE sequences positioned near each other in the genome undergo recombination after DNA damage, producing deletions, inversions, or other rearrangements. Using structural variant calls generated by the GraffiTE pipeline with Sniffles2, we focused on deletion- and inversion-mediated TEMRs.

From 36,790 deletions and 28,564 inversion structural variants identified across both cohorts, we detected 3,764 and 2,729 TEMRs, respectively (Supplementary Table 3). These events therefore accounted for ∼10.2% of all structural variants in the HBCC, and ∼9.6% in the NABEC, and most TEMRs were rare (HBCC=73.0%, NABEC=71%) (Fig. 5A), with many being singletons (HBCC=33.4% and NABEC=36.1%). Nearly all TEMRs were deletions (∼99% in both cohorts; Supplementary Table 3-4), consisting of 2,550 and 1,784 *Alu-Alu* pairs, 662 and 486 LINE-1-LINE-1 pairs, and 21 and 23 SVA-SVA pairs in HBCC and NABEC, respectively. TEMRs frequently overlapped with genic regions (64.8% in HBCC and 65.3% in NABEC), with notable differences in genomic localization: the majority of events in HBCC were exonic (63.4%), whereas in NABEC, TEMRs were more often intronic (50.4%).

**Figure 5.**
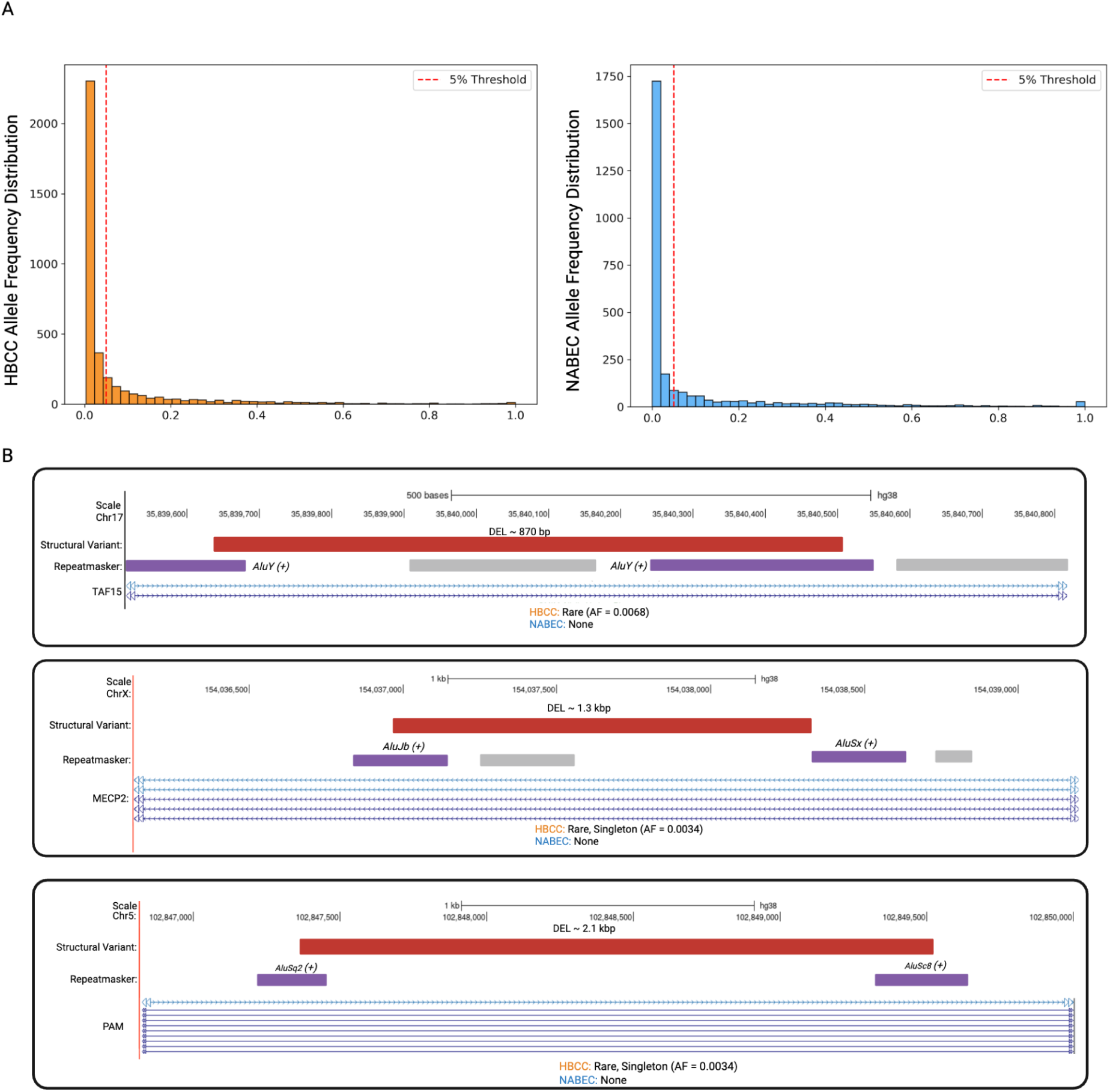
TE sequences mediate genomic rearrangements in the human genome. (A) Allele frequency distribution of TEMR events across both cohorts. The red line represents a 5% AF, where individuals at or below the red line are rare, and those above it are common. (B) Examples of TEMR Events that overlap with *TAF15* (top), *MECP2* (middle), and *PAM* (bottom). Schematic shows the structural variant (red bar) and flanking TE sequences (purple bars), along with the overlapping gene (bottom).

Several TEMRs overlapped genes associated with neurodevelopment and neurodegeneration. In the HBCC, we identified an 870 bp same-orientation *AluY–AluY*–mediated deletion, spanning intron 11 of *TAF15*, a gene implicated in frontal temporal dementia (FTD) (*33*) and ALS (*34*) (Fig. 5B). This rare heterozygous deletion was present in two individuals of African ancestry in HBCC, and absent in the NABEC, suggesting potential population specificity. We also identified a singleton 2.1 kb *AluJb–AluSx* deletion, in the same orientation, overlapping intron 1 of *PAM*, a gene linked to AD (*35*) (Fig. 5B), and a 1.3 kb *AluSq2–AluSc8* same orientation deletion, overlapping intron 2 of *MECP2*, a critical neuronal gene associated with Rett syndrome, a neurodevelopmental disorder (*36*) (Fig. 5B). This *MECP2* heterozygous deletion was observed in a single individual in the HBCC cohort.

### Transposable Elements Influence Gene Expression in the Frontal Cortex

To investigate the regulatory role of TEs in the human brain, we performed a TE-only eQTL (TE-eQTLs) analysis using TensorQTL on frontal cortex bulk short-read RNA-seq data from 205 NABEC and 76 HBCC individuals, using presence/absence genotypes of polymorphic TE insertions. From the GraffiTE TE calls, we retained only common (MAF > 5%) biallelic reference and non-reference TEs, pruning variants in LD (r² > 0.3) within a 1 Mb window to avoid redundancy. This yielded 2,564 TEs (1,709 reference; 855 non-reference) in HBCC and 1,877 TEs (1,309 reference; 568 non-reference) in NABEC.

At FDR q < 0.05, we identified 37 significant TE-eQTLs in HBCC (27 reference; 10 non-reference) and 196 in NABEC (148 reference; 48 non-reference), corresponding to 37 unique eGenes across 30 unique TEs in HBCC and 196 unique eGenes across 129 unique variants in NABEC (Supplementary Table 5-6). *Alu* elements accounted for the majority of significant TE-eQTLs, followed by LINE-1s and SVAs. Reference TEs showed an approximately equal distribution of positive and negative expression effects, while among non-reference TEs, SVAs showed a slight tendency towards increased expression and LINE-1s towards decreased expression (Supplementary Fig. 2). Together, these findings highlight TEs as both positive and negative regulators of gene expression in the frontal cortex, with some showing evidence of causal regulatory effects.

To compare TE regulatory effects with those of small variants, we performed a joint eQTL analysis (TE-joint-eQTLs), incorporating the same TEs with nearby SNVs and indels (Illumina-derived for NABEC = 257,822; ONT-derived for HBCC = 682,016). We identified 812 significant variant–gene associations in HBCC and 2,368 in NABEC, which included 811 unique eGenes (to 791 variants) in HBCC, and 2,238 unique eGenes (to 2,280 variants) in NABEC. Of these, TEs were the lead regulatory variant in two associations (both reference) for HBCC and in four associations (two reference and two non-reference) for NABEC (Supplementary Table 5-6). As expected, small variants showed bidirectional expression effects, while LINE-1 elements displayed opposite directional effects between reference and non-reference insertions (Supplementary Fig. 2).

To further prioritize TEs as causal variants relative to co-localising small variants, we applied CAVIAR, a Bayesian fine-mapping approach. TEs were identified as the most probable causal variant in five independent eQTL signals (HBCC: 2; NABEC: 3) (Supplementary Table 5-6). The most notable example was a 319 bp non-reference *AluY* insertion located 73.8 kb upstream of *CNTNAP3B* (q = 4.53×10^−6^; AF = 0.11), a gene previously associated with autism (*37*), where the insertion was the highest-probability causal variant associated with a decrease in *CNTNAP3B* expression (Fig. 6).

**Figure 6.**
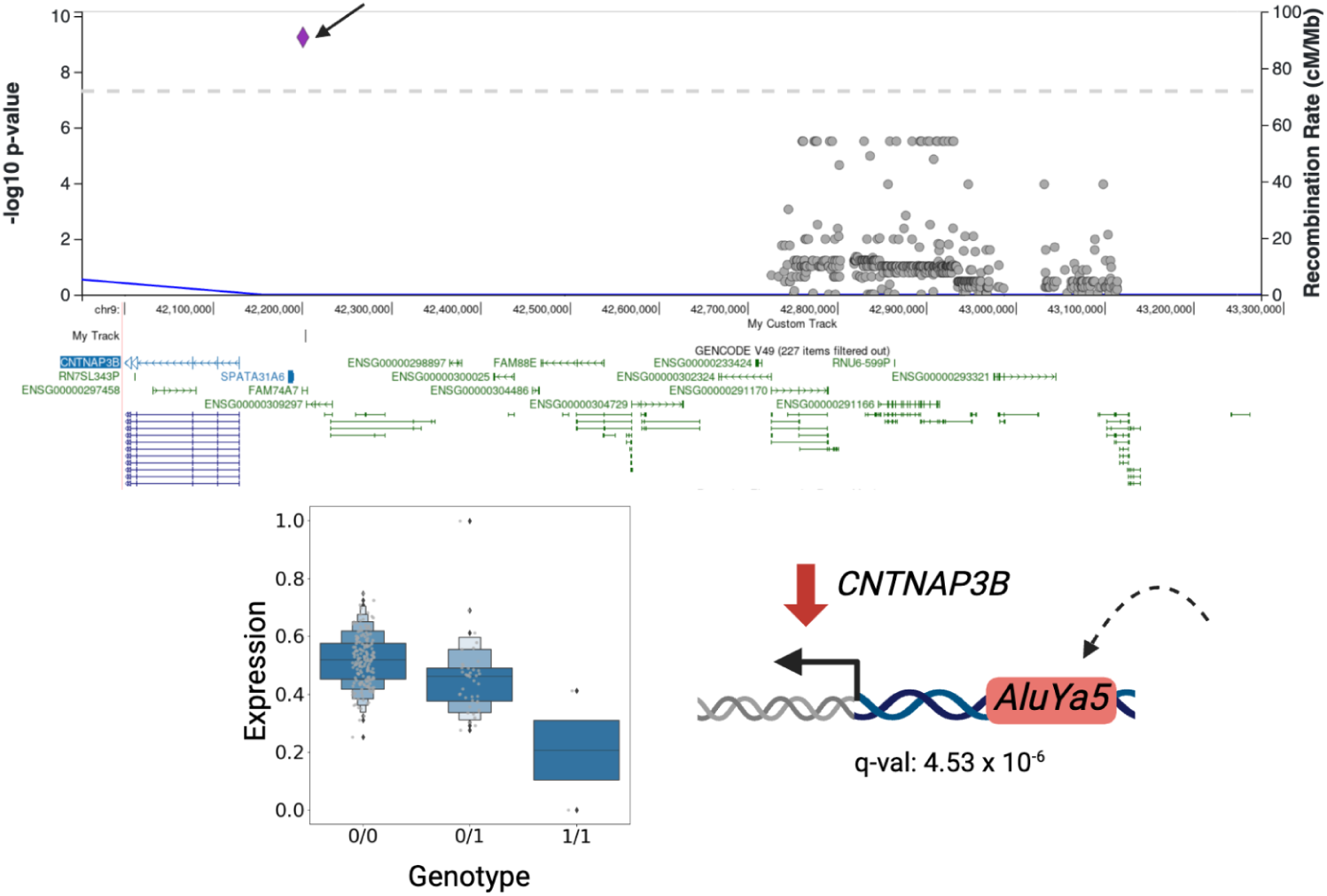
TE-eQTL and CAVIAR fine-mapping results for short-read bulk RNA-seq in NABEC. *Top*: LocusZoom plot showing the lead TE (purple diamond) and nearby small variants (grey circles) from the joint TE–small variant eQTL analysis. *Middle*: Genomic view of the TE, located 73.8 kb away from *CNTNAP3B*. *Bottom:* Boxplot of *CNTNAP3B* RNA expression by genotype, with accompanying schematic illustrating the potential impact of the TE on gene regulation.

### Transposable elements modulate gene expression in specific brain cell types

To investigate the cell-type-specific regulatory roles of TEs in the human brain, we performed single-nucleus TE-eQTL (TE-sn-eQTL) analysis across seven major frontal cortex cell types, applying the same TE filtering criteria as the bulk analysis (HBCC: 2,564 TEs; NABEC: 1,877 TEs).

Significant TE-gene associations were identified in both cohorts across multiple cell types. In HBCC, we detected 28 associations (24 reference; 4 non-reference) spanning astrocytes, excitatory and inhibitory neurons, oligodendrocytes, and oligodendrocyte precursor cells (Supplementary Table 7-8). A single reference *Alu* was associated with *KANSL1* and *LRRC37A2*, both implicated in AD and PD (*38*), across all of these cell types, indicating a shared regulatory signal rather than a cell-type-restricted effect. In the NABEC, 115 associations were identified (97 reference; 18 non-reference) across all seven cell types (Supplementary Table 7-8). In both cohorts, associations were enriched in glial lineages, with oligodendrocytes showing the highest number of significant associations, consistent with a widespread influence of TEs across brain cell populations rather than neuron-specific regulation.

A joint analysis incorporating TEs alongside small variants identified TEs as the lead regulatory variant in five associations in NABEC, all supported by CAVIAR fine-mapping as the most probable causal variants. These comprised three unique reference TE deletions: two in microglia and one in oligodendrocytes (Supplementary Table 7-8). A notable example was a 250 bp reference *Alu*Y deletion overlapping intron 5 of *SLC2A9* in microglia (q-value = 0.035; AF = 0.51), where the presence of the deletion allele was significantly associated with altered *SLC2A9* expression (Fig. 7). *SLC2A9* encodes the urate transporter GLUT9, previously implicated in PD through p53-mediated upregulation and modulation of intracellular urate levels (*39*).

**Figure 7.**
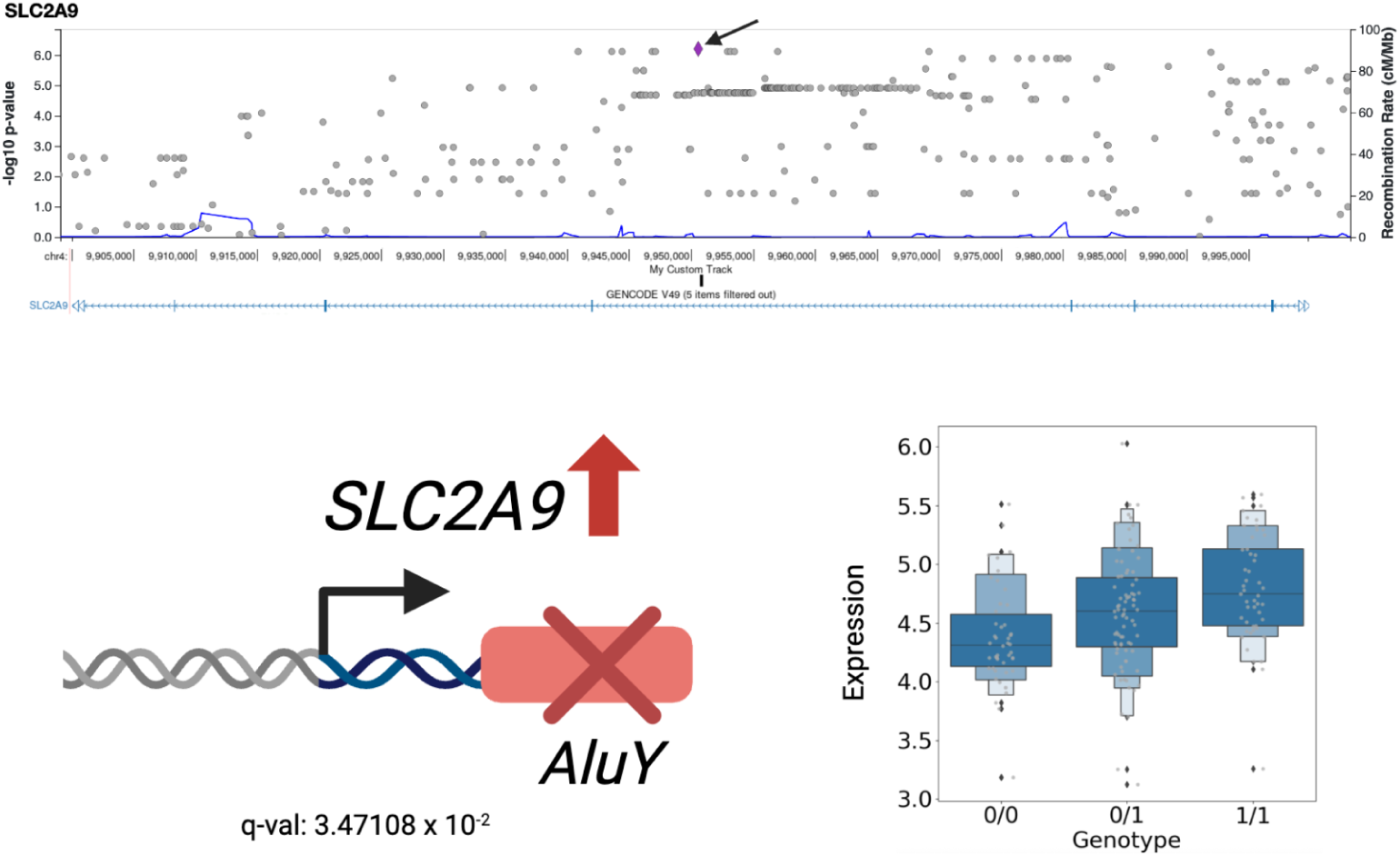
Single-nucleus TE-QTL and CAVIAR fine-mapping results for short-read snRNA-seq expression in NABEC. *Top*: LocusZoom plot showing the lead TE (purple diamond) and nearby small variants (grey circles) from the joint TE–small variant sn-QTL analysis. *Middle*: Genomic view of the TE, located within intron 5 of *SLC2A9*. *Bottom left*: Schematic illustrating the potential regulatory effect of the *AluY* deletion on *SLC2A9*. *Bottom right*: Boxplot of *SLC2A9* expression by genotype.

Directional effects varied by TE class and reference status across cell types. Small variants and non-reference *Alu* elements showed broadly symmetric, bimodal distributions of effect across most cell types. Reference *Alu* elements were consistently associated with higher expression across almost all cell types. Non-reference LINE-1 and SVA elements displayed more cell-type-specific directionality, with opposing effects observed between glial and neuronal populations (Supplementary Fig. 3).

### Transposable Element variants drive methylation differences across promoters, CpG islands, and gene bodies

To assess whether TE presence or absence is associated with local DNA methylation variation, we performed mQTL analyses across CpG islands (CGIs), gene bodies, and promoter regions, using the same TE filtering parameters applied in the eQTL and sn-eQTL analyses. This yielded 2,564 TEs (1,709 reference; 855 non-reference) in HBCC and 1,877 TEs (1,309 reference; 568 non-reference) in NABEC.

In TE-only analyses, we identified widespread TE–methylation associations across all three genomic contexts. In NABEC, we detected 46 TE–CGI (30 reference; 16 non-reference), 66 TE–gene body (45 reference; 21 non-reference), and 63 TE–promoter (38 reference; 25 non-reference) associations. In HBCC, we detected 14 TE–CGI (12 reference; 2 non-reference), 11 TE–gene body (5 reference; 6 non-reference), and 18 TE–promoter (12 reference; 6 non-reference) associations (Supplementary Tables 9–10). In joint analyses incorporating both TEs and small variants, association counts expanded across all contexts: in HBCC, we identified 4 CGI, 8 gene body, and 8 promoter associations; in NABEC, 47 CGI, 75 gene body, and 8 promoter associations. Fine-mapping with CAVIAR identified TEs as the most probable causal variant at a subset of these loci, including 3 CGI and 5 gene body mQTLs in NABEC, and 1 promoter mQTL in HBCC, providing evidence that TEs themselves, rather than co-localising small variants, drive local methylation changes.

A notable example is a 325 bp non-reference *AluY* insertion (AF = 0.39) in NABEC, which was the top-ranked causal variant at two independent CGI associations with opposing directions of effect (Fig. 8). Despite proximity to multiple annotated gene loci, this insertion showed no corresponding signal in eQTL analyses, suggesting its regulatory influence operates primarily at the level of DNA methylation rather than steady-state transcription.

**Figure 8.**
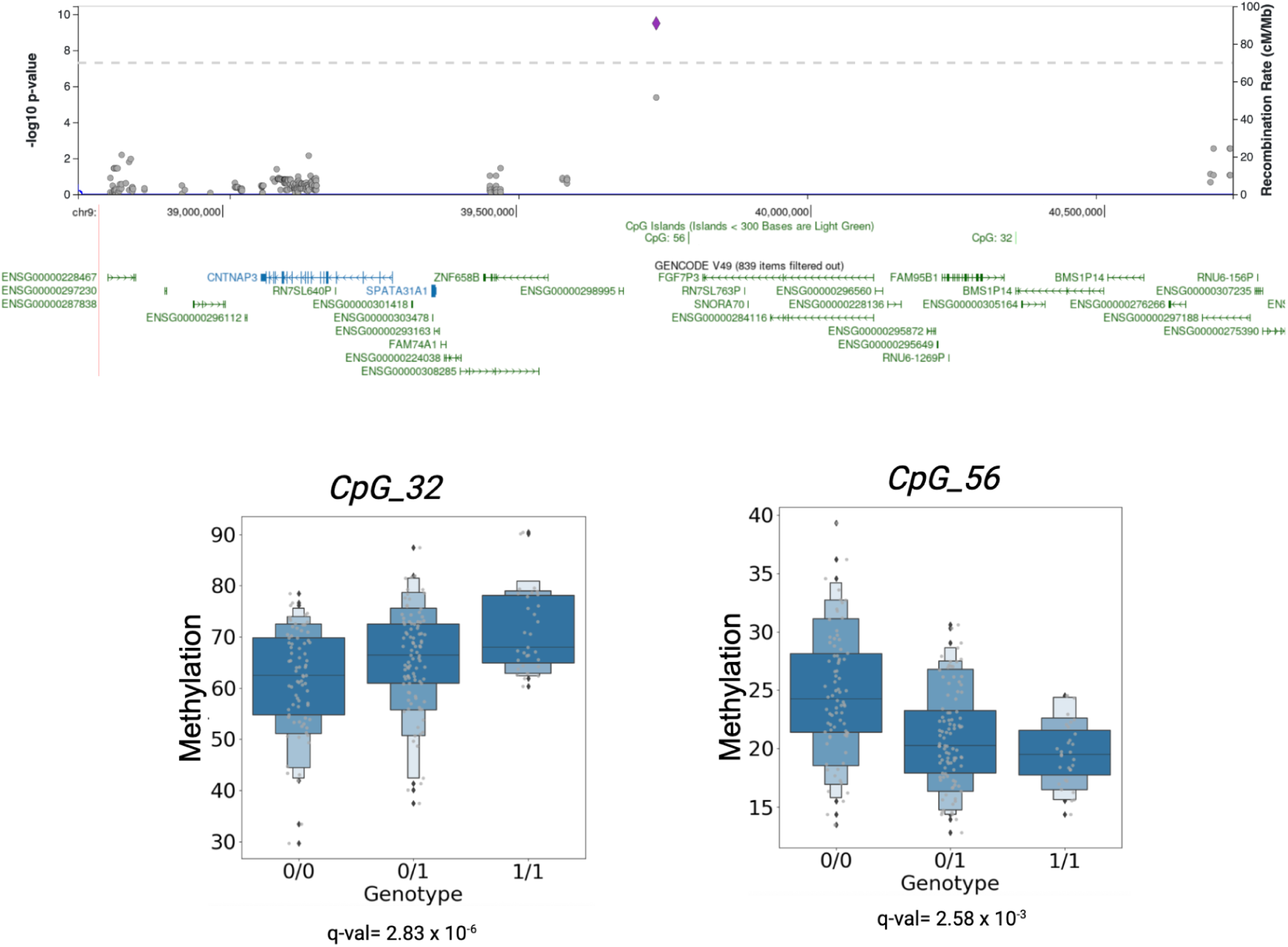
Methylation QTL and CAVIAR fine-mapping results from long-read sequencing methylation data in HBCC. *Top*: LocusZoom plot showing the lead TE (purple diamond) and nearby small variants (grey circles) from the joint TE–small variant mQTL analysis. *Middle*: Genomic view of the TE located between two CpG islands: CpG_56 and CpG_32. *Bottom*: Boxplot showing methylation levels stratified by genotype for each.

Across both cohorts, TE-mQTLs showed broadly balanced directional effects (bimodal beta-value distributions); however, non-reference *Alu* elements were consistently associated with higher methylation at nearby gene bodies, while reference *Alu* deletions showed a bias towards reduced methylation (Supplementary Fig. 4). This pattern is consistent with a model in which recently inserted, non-reference TEs are subject to stronger epigenetic silencing that spreads into flanking regions, whereas older, reference-annotated elements are less actively repressed and may instead contribute permissive chromatin environments at nearby regulatory sequences.

### Global TE Hypermethylation and Age-Linked *Alu* Demethylation in the Frontal Cortex

Whereas the mQTL analyses above assessed how TE variation influences methylation at flanking genomic loci, we next sought to characterise the methylation state of TE sequences themselves. To do so, we analyzed haplotype-resolved CpG methylation aggregated directly across reference TE bodies in frontal cortex samples from both cohorts, enabling a population-scale view of the epigenetic landscape within each TE family. Consistent with prior reports of TE silencing in long-read CpG datasets, *Alu*, LINE-1, and SVA elements were broadly hypermethylated, with ≥80% CpG methylation across most loci (Supplementary Fig. 5).

Subfamily-level analyses revealed that *Alu* elements maintained hypermethylation across all evolutionary lineages *(AluJ, AluS, AluY*, and monomeric), although monomeric *Alus*, considered the oldest, displayed modest hypomethylation in both cohorts (Supplementary Fig. 6), consistent with gradual methylation loss over evolutionary time due to sequence divergence or reduced epigenetic repression. LINE-1 elements were uniformly hypermethylated across subtypes (Supplementary Fig. 7). For SVAs, most loci were hypermethylated, supporting the hypothesis that newly inserted elements undergo epigenetic silencing to maintain transcriptional stability (*40*). SVA elements are classified into six subfamilies (SVA_A through SVA_F) based on sequence divergence, with SVA_A representing the oldest and SVA_F the youngest and most human-specific. Most SVA loci were hypermethylated across all subfamilies, supporting the hypothesis that newly inserted elements undergo epigenetic silencing to maintain transcriptional stability. However, when grouping by SVA class, subsets of SVA_A and SVA_E (and SVA_C in NABEC) exhibited apparent hypomethylation (Supplementary Fig. 8); these loci were <500 bp and likely represent truncated fragments rather than full-length SVAs, suggesting their hypomethylation has limited biological relevance.

We next tested whether TE methylation correlated with chronological age. *Alu* elements were the only TE class to display consistent age-associated methylation changes across both cohorts, with 12 significant loci in HBCC and 3,504 in NABEC across both haplotypes. However, the significant difference in the number of associations between the two cohorts may be due to the wider age range in NABEC (15–96 years vs. 18–85 in HBCC) and greater representation of older individuals (Supplementary Table 1). Although 44% of age-associated *Alu*s showed hypermethylation with age, the majority (56%) exhibited hypomethylation (Fig. 9). Most age-associated *Alu*s overlapped intronic regions (69.0% in NABEC; 66.7% in HBCC), and Gene Ontology enrichment of overlapping genes in NABEC highlighted pathways involved in neurogenesis, neuronal differentiation, and neuron development (Supplementary Fig. 8).

**Figure 9.**
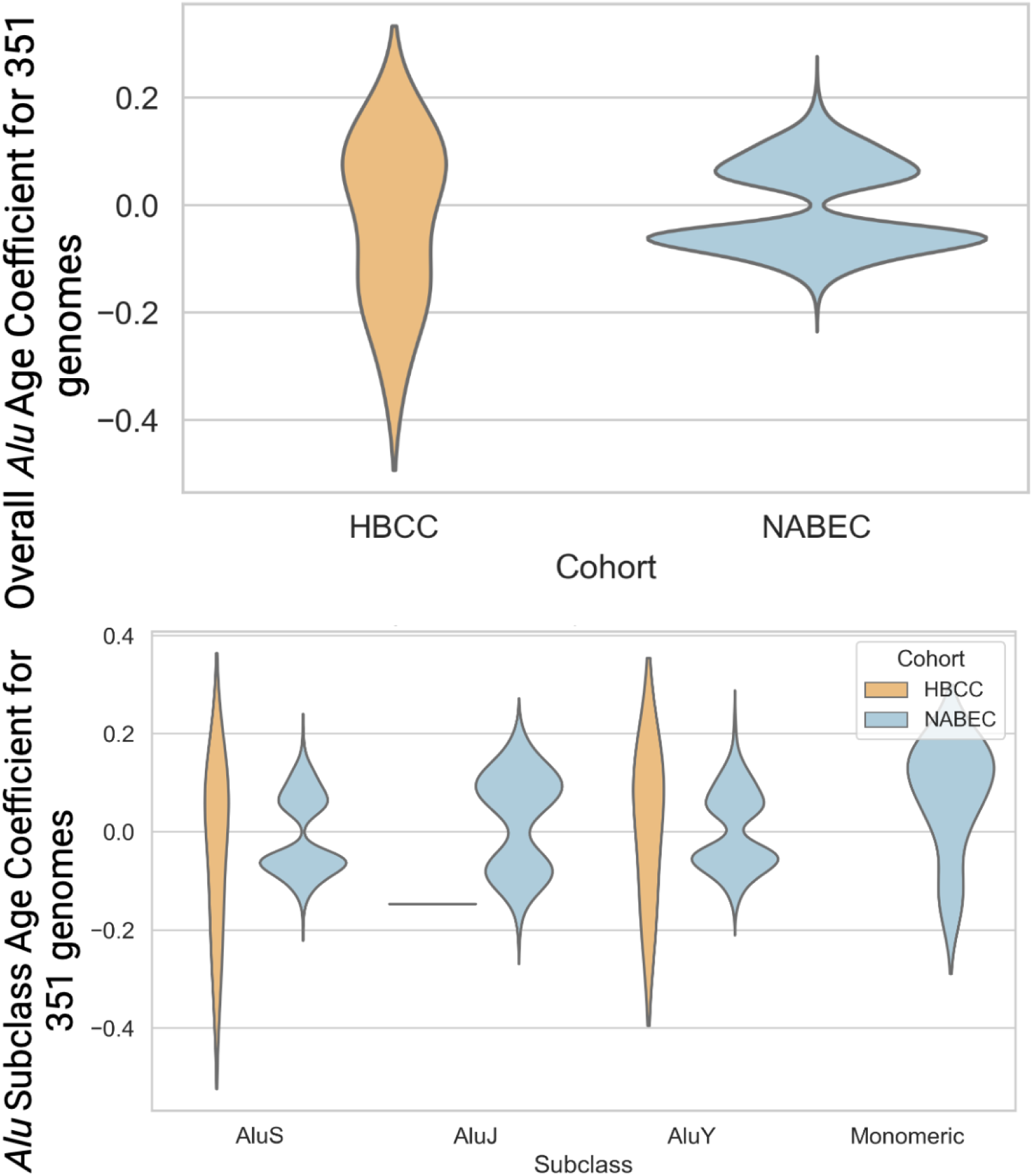
Linear age regression analysis of TE methylation across diverse populations. *Top*: Overall methylation changes with age for both haplotypes, combined, in HBCC and NABEC. *Bottom*: *Alu* subclass specific methylation changes with age for both haplotypes combined, in HBCC and NABEC. In both cohorts, most *Alus* exhibit hypomethylation, with subclass-specific differences in the magnitude and direction of change.

## Discussion

In this study, we leveraged long-read whole-genome sequencing to generate a comprehensive population-scale catalogue of germline transposable element variation in the human brain, analysed across 351 postmortem frontal cortex samples from two ancestrally diverse cohorts. By integrating TE detection with structural variant analysis, expression and methylation QTL mapping, single-nucleus transcriptomics, and direct long-read CpG profiling, we demonstrate that TEs are widespread contributors to genetic and epigenetic variation in the human brain, roles that have been largely inaccessible to short-read approaches.

Using GraffiTE, we identified over seven thousand high-confidence polymorphic TEs, with more than half representing non-reference insertions, many of which were rare or population-specific. As expected, the majority of insertions were *Alu*s, followed by LINE-1s and SVAs, and were most frequently located within intronic regions, consistent with prior reports of TE distribution across the genome (*26*). The HBCC cohort, comprising individuals of African and African-admixed ancestry, harboured a greater number of TE insertions compared to NABEC, consistent with the higher levels of genetic diversity documented in African-descent populations (*41*).

Beyond presence/absence detection, long-read sequencing enabled us to characterise the internal architecture of reference SVA and *Alu* elements at individual loci, revealing that the majority are multiallelic, with substantial diversity in both motif sequence and repeat length. This level of structural complexity is largely inaccessible to standard variant calling approaches and has not previously been reported at population scale in brain tissue. At the *KANSL1* locus, an intronic STR within a reference *AluY* element and an independent SVA_D VNTR were each independently associated with increased gene expression, demonstrating that repeat length variation within distinct TE families can independently modulate expression of the same gene. The same effect was found on the single cell level, in which multiple cell types were significantly associated with changes in expression at the same locus in the sn-QTL analysis. This finding is further supported by experimental evidence from Fröhlich et al., who demonstrated that CRISPR-mediated deletion of a separate SVA element at the same locus directly increases *KANSL1* expression, a result independently corroborated here using a distinct reference SVA (*42*). Notably, certain highly complex loci cannot be fully resolved even with long-read sequencing; the *MAPT* inversion region, owing to its exceptional structural complexity, remains beyond the resolution of current long-read approaches, and almost certainly harbours additional repeat-mediated variation with functional consequences for local gene expression and methylation.

We also identified a substantial number of TEMRs with approximately 10% of structural variants across both cohorts arising through homology-mediated recombination between flanking TE sequences, many of which were deletions. TEMRs were more frequent in HBCC than in NABEC, paralleling the overall higher levels of SV and TE variation observed in that cohort. Most events were rare, including singletons and ancestry-specific variants, and the majority were *Alu–Alu* recombination events, consistent with the relative abundance of *Alu* elements in the human genome. Several TEMRs overlapped genes implicated in brain function and neurodegenerative disease, including a 2.1 kb singleton deletion in HBCC overlapping *PAM*, a gene previously linked to AD (*35*). The functional consequences of TEMRs have been demonstrated at specific disease-relevant loci; intronic *Alu* elements can drive premature transcript termination, alter A-to-I RNA editing, and generate alternative isoforms through secondary structure formation, mechanisms with clear relevance to the neurodegenerative disease genes implicated among our TEMR calls (*43*). Together, these findings demonstrate that TEs contribute to rare, population-specific structural variation and expand the landscape of genomic diversity in ways that may influence neurodegenerative disease risk.

We next examined the regulatory impact of TEs through eQTL analyses, leveraging both bulk tissue and cell-type-resolved transcriptomic profiles. Using TensorQTL, we identified numerous TE–gene association pairs, with a greater number detected in NABEC compared to HBCC, most likely reflecting differences in sample size and transcriptomic coverage rather than ancestry-specific effects. Integrating TE calls with small-variant data, we identified multiple instances in which TEs were the top-ranked candidate causal variants driving gene expression. Extending these analyses to single-nucleus data, we identified a subset of TE–gene associations in which fine-mapping pointed to TEs as the most likely causal variants within specific cell types, most prominently oligodendrocytes and microglia. TEs are well-established contributors to cis-regulatory architecture, providing promoters, enhancers, and transcription start sites, and genic TE insertions that can alter splicing and transcript structure to influence expression levels (*44–46*). As the eQTL and sn-eQTL frameworks are primarily powered to detect common variants in neurologically healthy individuals, the TE-driven regulatory effects we identify are unlikely to reflect deleterious consequences; rather, they highlight how TE insertions can be assimilated into cell-type-specific regulatory programmes in ways that are tolerated or co-opted over evolutionary time.

We further demonstrate that TEs contribute to epigenetic regulation through mQTL analyses, with TE-associated methylation effects widespread across gene bodies, CGIs, and promoters; though the most pronounced effects were observed within gene bodies. Fine-mapping frequently identified the TE itself as the most likely causal variant. Notably, reference deletions and non-reference insertions exhibited distinct behaviours within *Alu* elements in gene bodies, with non-reference insertions consistently associated with increased local methylation, consistent with a model epigenetic silencing of newly integrated elements extends into flanking sequences, reshaping local chromatin states and influencing the accessibility of nearby regulatory elements, though it is plausible that some of these TEs are simply adopting the pre-existing methylation state of their surroundings (*47*). In contrast, hypomethylation was observed predominantly at reference TE loci, consistent with the notion that older elements, having persisted over evolutionary timescales, may require less active silencing and can instead be repurposed as regulatory elements, with reduced methylation potentially enabling enhancer-like activity at proximal genes (*48*).

To complement the mQTL analysis, we profiled CpG methylation across TE sequences at single-base resolution using long-read sequencing, enabling simultaneous detection of TE variation and methylation with a level of locus-specific precision not achievable with bisulfite or short-read approaches. Consistent with previous literature, TEs were globally hypermethylated in the frontal cortex, reflecting robust epigenetic silencing of repetitive elements in adult brain tissue (3, 8). We additionally observed modest hypomethylation in evolutionarily older *Alu* subfamilies, suggesting that methylation maintenance at these elements may decrease gradually over time, likely as a consequence of accumulated sequence divergence (*40*). Among the TE families examined, only *Alu* elements displayed pronounced age-associated methylation changes, with progressive demethylation enriched near genes implicated in neuronal differentiation and neurogenesis, consistent with prior reports of associations between decreased *Alu* methylation and age (*49*), and suggesting functional relevance to brain aging. No significant age-associated methylation changes were identified for LINE-1s or SVAs. Loss of methylation at *Alu* elements may increase their transcriptional activity and propensity for exonisation, a phenomenon previously documented in the frontal cortex, where it can influence alternative splicing and transcript structure. While exonisation has been proposed as a source of transcriptomic diversification, it may also generate aberrant isoforms with potential relevance to neurodegeneration. Future studies leveraging long-read RNA sequencing will be essential to directly resolve these events and distinguish their contributions to normal brain aging from those associated with pathology.

Several technical considerations shape the interpretation of these findings. Differences in sequencing chemistry between cohorts, R9.4.1 for NABEC and R10 for HBCC, may affect variant detection, particularly at multiallelic loci. The use of short-read RNA sequencing for QTL analyses limits isoform-level resolution, which is especially relevant given the established role of TEs in alternative splicing. Our biallelic variant framework excludes multiallelic TE alleles that are common in repetitive regions, and reliance on GRCh38 restricts characterisation of centromeric and pericentromeric TE variation. Future studies incorporating telomere-to-telomere assemblies, long-read transcriptomics, and somatic TE detection will be important for extending this work, as will applying this framework to disease cohorts to directly examine the contribution of TEs to neurodegeneration risk.

In summary, this study provides the most comprehensive population-scale characterisation of germline TE variation in the human brain to date, demonstrating that TEs are widespread contributors to structural variation, gene regulation, and age-associated epigenetic change. We release a publicly available catalogue of TE variation, including allele frequencies stratified by ancestry across European and African/African-admixed populations, as a resource for the neurogenomics community, providing a foundation for future population-genetic, functional, and disease-focused investigations.

## Methods

### Long-read Data Sources and Prior Processing

We analyzed publicly available ONT long-read sequencing data from postmortem human brain samples previously generated by Billingsley et al. (*27*). These datasets were accessed through the NIH AnVIL platform and the Alzheimer’s Disease Workbench (ADWB), and comprise neurologically healthy control samples from two cohorts: the North American Brain Expression Consortium (NABEC) and the Human Brain Collection Core (HBCC).

The NABEC cohort includes frozen frontal cortex tissue from 205 individuals of European ancestry with no clinical history of neurological disease. The HBCC cohort includes frontal cortex tissue from 146 individuals of African and African admixed ancestry. Full cohort metadata, including age at death and sex distribution, are available on AnVIL and ADWB and summarized in Supplementary Table 1.

Tissue collection for NABEC was approved by the Joint Addiction, Aging, and Mental Health Data Access Committee (dbGaP accession phs001300.v4.p1). HBCC samples were collected under CNS IRB–approved protocols (NCT03092687), with next-of-kin consent obtained via the Offices of the Chief Medical Examiners in the District of Columbia and Virginia. Per NIH policy, the use of postmortem tissue qualifies as non–human subjects research.

Sequencing protocols are publicly available on protocols.io (*50*) and described in detail in Billingsley et. al. (*27*). In brief, 40 mg of frozen brain tissue was homogenized, and high-molecular-weight DNA was extracted and sheared to ∼30 kb. NABEC libraries were prepared using the ONT SQK-LSK110 kit and sequenced on PromethION R9.4.1 flow cells (FLO-PRO002), while HBCC libraries were prepared with the SQK-LSK114 kit and sequenced on R10.4.1 flow cells.

Basecalling was previously performed using Guppy v6.1.2 for NABEC and v6.3.8 for HBCC. All samples were processed using the NAPU pipeline, which produced BAM files aligned to the GRCh38 reference genome via minimap2 v2.23-r1111 (*51*), and generated small variant (SNV/indel), structural variant (SV), and DNA methylation calls. Small variants (<50 bp) for HBCC were called using DeepVariant with the ONT_R104 model. Due to lower basecalling accuracy for SNVs from R9 flow cells, short-read Illumina WGS data (dbGaP accession phs002636.v3.p1) was used for NABEC to supplement variant calling. Additional Illumina data for NABEC was accessed via dbGaP accession phs002636.v3.p1.

### Transposable element and structural variant calling

Structural variants were identified using the GraffiTE (*28*), which employs Sniffles2 (v2.6.3) to detect reference-based, unphased structural variants from long-read BAM files. Only insertions and deletions were retained for downstream analysis. To annotate TEs, we utilized a custom TE library derived from Dfam (https://dfam.org/releases/current/families/Dfam-RepeatMasker.lib.gz), which was converted into FASTA format using the FAMDB tool (v2.0.2). This FASTA was provided as input to RepeatMasker (v4.1.3), and annotations were filtered to retain only TE insertions spanning ≥80% of the corresponding SV length. To improve genotyping accuracy and capture complex variation, presence/absence genotypes were recalculated using GraphAligner (v1.0.20) and vg (1.54.0). For this, a genome graph was constructed with TEs represented as bubbles, enabling more robust alignment and variant resolution. TEs were classified as reference or non-reference based on overlap with the UCSC RepeatMasker track. SV-associated TEs overlapping UCSC RepeatMasker annotations were designated as reference, while those lacking such overlap were classified as non-reference or novel.

### Repeat calling

To characterize the tandem repeat loci within reference SVA and *Alu* TE annotations, a customized VAMOS motif file was made. Briefly, previously formulated VAMOS motifs (v2.1, https://github.com/ChaissonLab/vamos) were overlapped with Repeatmasker *Alu* and SVA reference sequences vis BEDTools intersect (v. 2.31.1), in which motifs which overlapped were kept. We ran VAMOS (v. 2.1.7) utilizing both those motifs and phased BAM long-read sequencing files for both NABEC and HBCC, previously generated by Billingsley et al. (*27*). From here, tryvamos (v. 1.2.0) was used to extract relevant repeat length data, and a simple linear regression (Expression ∼ VNTR repeat length + Age + Sex + PMI) was run.

### Expression datasets

#### Bulk short-read RNA sequencing data

To determine the impact of newly identified TEs, sample data was taken from the North American Brain Expression Consortium (dbGaP Accession phs001300.v4.p1), in which sample bulk RNA-seq data was quantified using Salmon with the filtered Gencode v43 human transcriptome index. The expression data featured 206 samples, with 60,880 quantified genes, in which sex - specific ones were subsetted and plotted to identify any sample gender mismatches. Any genes with a missingness over 0.25 were excluded, leaving 26,778. Data was then quantile transformed and scaled 0 to 1. Data was then normalized by quantile-normalization and min-max scaled. For the HBCC sample data was taken from the Human Brain Collection Core (https://nda.nih.gov/edit_collection.html?id=3151), and similar to NABEC, bulk RNA-seq data was quantified with Salmon with the filtered Gencode v32 transcriptome index. The sample count was 86 covering 60,880 quantified genes, in which sex specific genes were also subsetted and plotted to identify gender mismatches. Genes with a missingness over 0.25 were excluded, leaving 26,679 genes. Data was then normalized by quantile-normalization and min-max scaled.

### Single-Nucleus short-read multiome dataset

We utilized publicly available processed single-nucleus multiome (*52*) (ATAC and gene expression) data generated from postmortem dorsolateral prefrontal cortex (DLPFC) samples of neurologically healthy individuals of European and African admixed ancestry (NABEC data is available on dbGaP under accession phs004202.v1.p1; HBCC multiome data is available on the NIMH Data Archive (under accession 3151 as CARD PFC Multiome snRNA, https://nda.nih.gov/edit_collection.html?id=3151). This resource includes 202 individuals from the NABEC and 155 individuals from the HBCC. Across both cohorts, donors ranged in age from 15 to 100 years, with mean ages of 48.8 (NABEC) and 45.2 (HBCC) years. Single nuclei multiome profiling was performed on a total of 2.83 million nuclei, with stringent quality control filtering for ambient RNA, ATAC fragments, number of expressed genes, ribosomal and mitochondrial transcript content, and doublets, resulting in >1.5 million high-quality nuclei across seven major brain cell types (astrocytes, excitatory neurons, inhibitory neurons, microglia, oligodendrocytes, oligodendrocyte precursor cells, and vascular cells).

### Methylation dataset

#### Bulk long-read methylation data

Phased methylation data for both the NABEC and HBCC cohorts were previously generated, as described in Billingsley et. al.(*27*, *51*). Briefly, Modkit pileup was used to generate methylation BED files from ONT long-read data aligned to the GRCh38 reference genome. Methylation calling was restricted to 5-methylcytosine (5mC) at CpG dinucleotides, and calls were combined across strands to create strand-agnostic representations. Aggregated methylation profiles were generated using bedtools map, with regions filtered to retain only those containing a minimum of 10 CpG sites and at least 5 supporting reads. These aggregated values were then averaged across gene promoters, gene bodies, and CpG islands (CGIs). Promoter coordinates were obtained from the Eukaryotic Promoter Database and extended by 1 kb upstream and downstream of each transcription start site. Gene body regions were defined using the UCSC genome annotation (refFlat) for the GRCh38/hg38 assembly (Dec. 2013), selecting the transcript isoform with the greatest number of exons to represent each gene. CpG island coordinates were obtained from the UCSC Genome Browser CpG Island track. Final methylation values were normalized per site plus coverage prior to downstream analyses.

TE methylation profiles were derived from the same CpG methylation calls described above. Aggregated methylation values across TEs were calculated using bedtools, based on TE coordinates from the UCSC RepeatMasker track for GRCh38. Methylation levels were averaged across CpG sites within annotated TE regions, focusing on the three major TE families: *Alu,* SVA, and LINE-1. To ensure robustness, only TE instances with a minimum of 95% of reads having CpG sites were kept. TE loci with missing methylation data in more than 5% of samples were excluded from downstream analysis.

In order to compare an equal number of CpGs across samples, we then only compared CpGs that were present in all samples. We achieved this by removing individual CpGs that were deleted in any sample within each cohort, which was also extended CGIs, promoters, and enhancers. This allowed us to calculate average regional methylation across an equal number of CpGs in that gene body region.

### Statistical Analysis

#### Expression and Methylation QTL Mapping

Cis-expression and methylation quantitative trait locus (QTL) analyses were performed using previously generated gene expression and DNA methylation data with the TensorQTL framework. Analyses were conducted independently within each cohort, incorporating normalized expression or methylation levels as traits and genotypes as predictors. Covariates included sex, age, postmortem interval, dataset group (e.g., SH, UKY, or UMARY for NABEC), and principal components (PCs) derived from genotype and trait data. Genetic and trait PCs were generated using the PCA module from the *scikit-learn* Python package, and scree plots were used to retain components that collectively explained approximately 70% of the variance. Missing covariate data were imputed using median values via the *SimpleImputer* function from *scikit-learn*.

Genotype data were prepared using PLINK v1.9. Only autosomal, bi-allelic variants with a minor allele frequency greater than 0.05 were retained for downstream analysis. QTL testing was performed using both TE-only and combined TE + SNV genotype datasets. Associations were calculated for all variant–trait pairs located within ±1 Mb of the transcription start site of each gene or methylation feature. Multiple testing correction was performed using the q-value method, with a false discovery rate threshold of q ≤ 0.05 to define statistical significance. To explore the overall distribution of effect directions across variant types, we visualized slopes for eQTLs passing a nominal significance threshold based on pval_beta < 0.05, which reflects per-gene correction for the number of variants tested. This more inclusive threshold captures the broader landscape of directional effects without being limited to genome-wide significant associations.

#### Fine-Mapping of Causal variants

To identify putative causal variants among the significant QTL associations, we applied CAVIAR(*53*), a fine-mapping method that estimates the posterior probability of causality for each variant by integrating summary association statistics with local linkage disequilibrium (LD) patterns. For each gene or methylation site with a significant QTL (q ≤ 0.05), Z-scores were calculated from the effect size and standard error of the variant–trait associations. The top 100 variants, ranked by Z-score magnitude, were selected as input to CAVIAR. Analyses assumed a single causal variant per locus (causal set size = 1). LD matrices were derived from the corresponding genotype data, and CAVIAR posterior probabilities were used to prioritize candidate causal variants for both eQTLs and mQTLs. structural variants were selected as most probable causal variants when they had the highest posterior probability at a locus as well as when tied with an SNV for the highest probability.

#### Methylation Age Regression Analysis

To explore associations between TE methylation and chronological age, we performed linear regression analyses across all reference TE-associated methylation regions. For each region, methylation levels were modeled as the dependent variable and age was included as the primary predictor, with methylation principal components, postmortem interval, and sex included as covariates. Regression analyses were conducted separately within each cohort. Multiple testing correction was applied using the Benjamini–Hochberg method, with FDR-adjusted p-values ≤ 0.05 considered statistically significant.

#### Transposable Element Mediated Rearrangement (TEMR) Analysis

We evaluated the impact of TEs on structural variants using a pipeline generated by Balachandran, Parithi, et al. (*32*). TEMR events were determined using BEDTools intersect (*54*), in which SV data were used to assess breakpoint coordinates, extended by 2 bp on either side, which were then intersected with TE coordinates. Repeats were then placed at each side of the structural variant, depending on their genomic location. Any events with multiple repeats overlapping either breakpoint or with more than 80 percent of the SV spanned by a single TE family were excluded from the final TEMR events table. If two repeats (one at each end) were of the same subfamily, in the same or opposite orientations, the structural variant was flagged as a potential TEMR, then labelled accordingly. Using gene track information from UCSC Gencode V48, TEMR events were then overlapped using BEDTools Intersect (*54*) to determine gene proximity.

#### Pathway Analysis

We generated gene lists for NABEC and HBCC based on the results of the methylation-by-age regression analysis. Gene lists were created by identifying UCSC gene annotations (Gencode v48) that overlapped with significant regression hits, as well as a background gene set consisting of all genes that overlapped with any loci included in the regression analysis. Overlaps were determined using BEDTools Intersect (*54*). Using FUMAgwas (59), Gene Ontology (GO) enrichment for biological processes was performed and the results were then reported.

## Code availability

The code used to process and analyze the data for this study is publicly available at https://github.com/NIH-CARD/CARDlongread_NABEC_HBCC_TE_manuscript. This repository includes scripts and workflows for processing of GraffiTE TE annotation output, eQTL and mQTL analyses, and downstream data integration. Additional details about the computational tools and parameters used in this study are described in the Methods section of the manuscript.

## Data availability

Human brain sequencing datasets are under controlled access and require a dbGap application (phs001300.v4) (phs000979.v4). Afterwards, the data will be available through the restricted AnVIL workspace. All other data and code needed to evaluate and reproduce the results in the paper are present in the paper and/or the Supplementary Materials.

## Supplementary Figure Legends

**Supplementary Figure 1.**
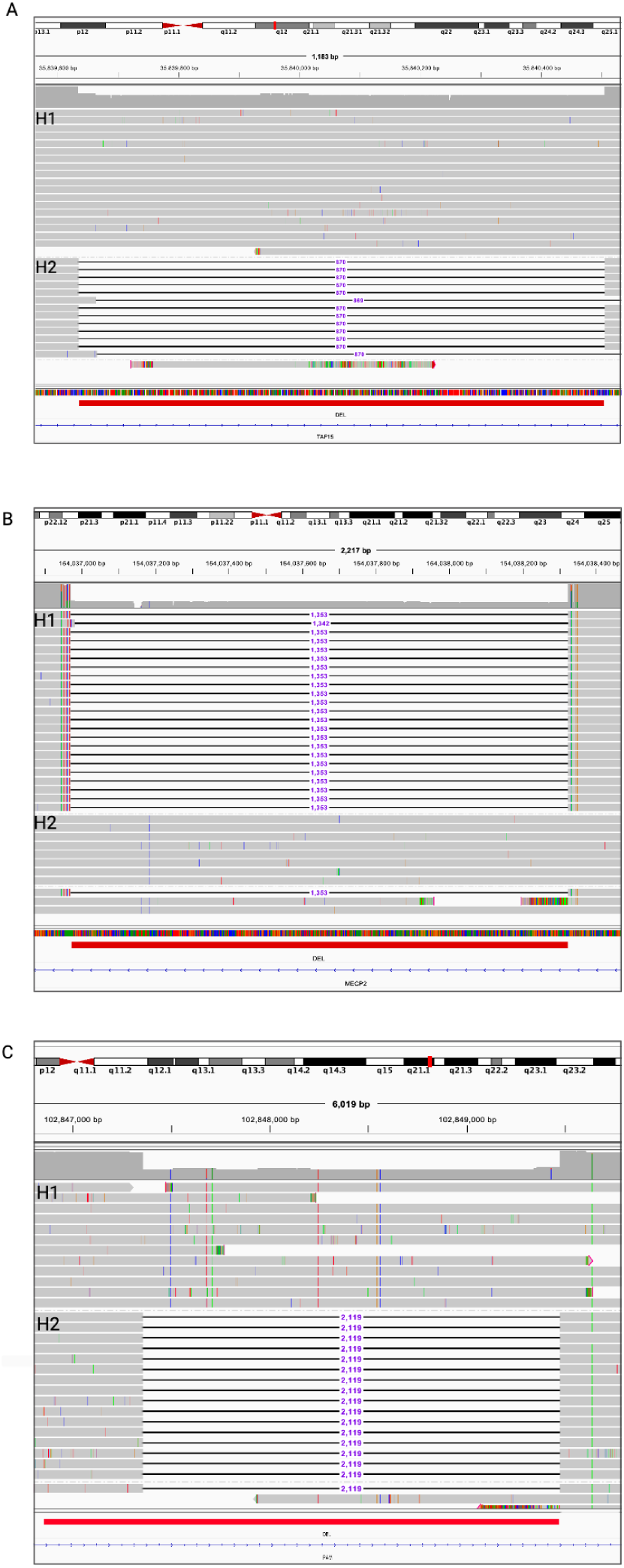
IGV Validations of TEMRs across Structural Variant Carriers. TEMR event examples in HBCC, which overlap with (a) *TAF15*, (b) *MECP2*, and (c) *PAM* loci, were validated in IGV as heterozygous deletions within carrier samples.

**Supplementary Figure 2.**
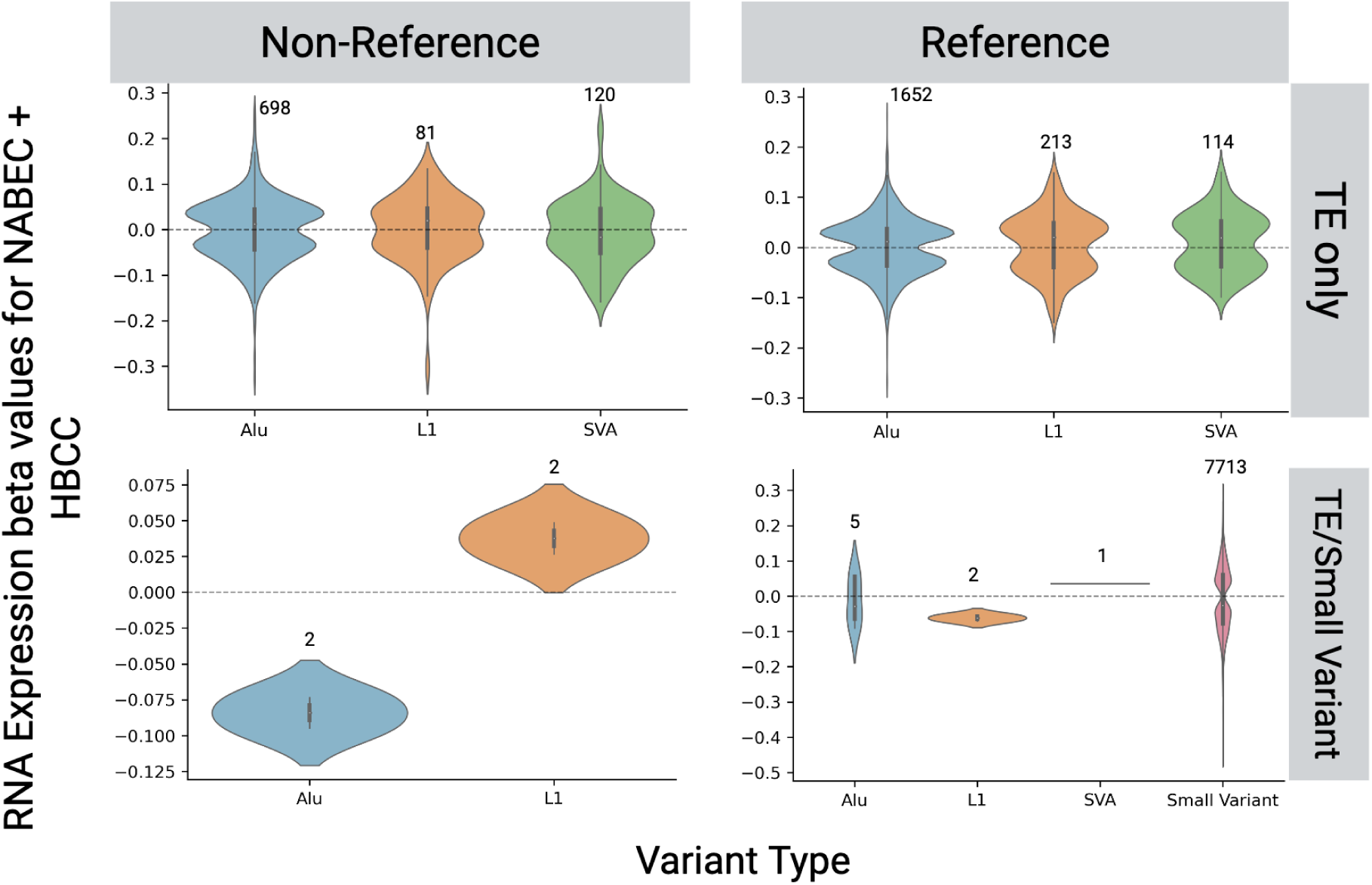
Comparison of RNA expression effects by variant types. *Top Left*: TE-only eQTL by TE type for non-reference (insertions). *Top Right*: TE-only eQTL by TE type for reference (deletions). *Bottom left*: TE-Small variant joint eQTL by TE type for non-reference (insertions). *Bottom Right*: TE-Small variant joint eQTL by TE type for reference (deletions).

**Supplementary Figure 3.**
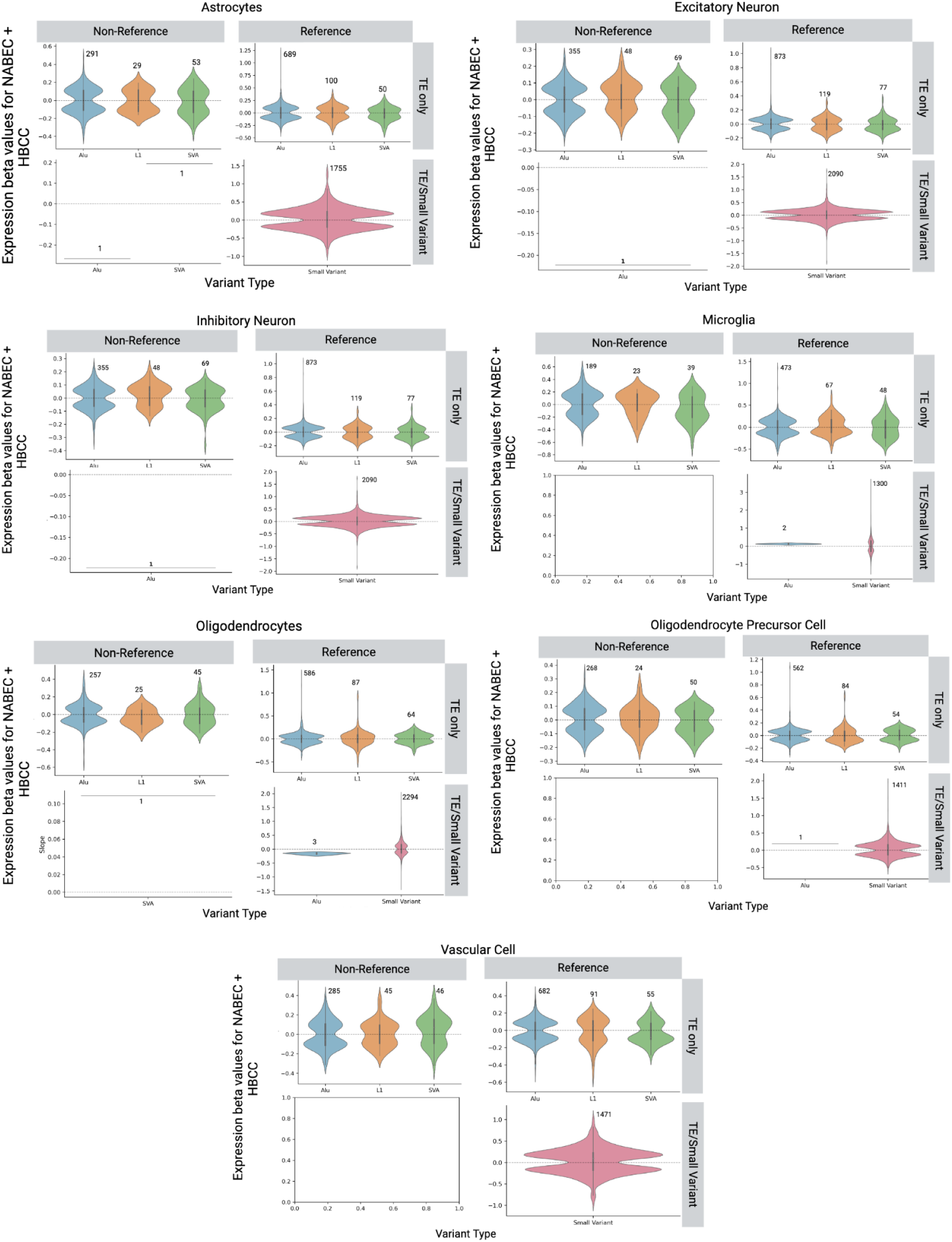
Comparison of single-nucleus RNA seq expression by 7 cell types. For each panel: *Top Left*: TE-only sn-QTL by TE type for non-reference (insertions). *Top Right*: TE-only sn-QTL by TE type for reference (deletions). *Bottom left*: TE-Small variant joint sn-QTL by TE type for non-reference (insertions). *Bottom Right*: TE-Small variant joint sn-QTL by TE type for reference (deletions).

**Supplementary Figure 4.**
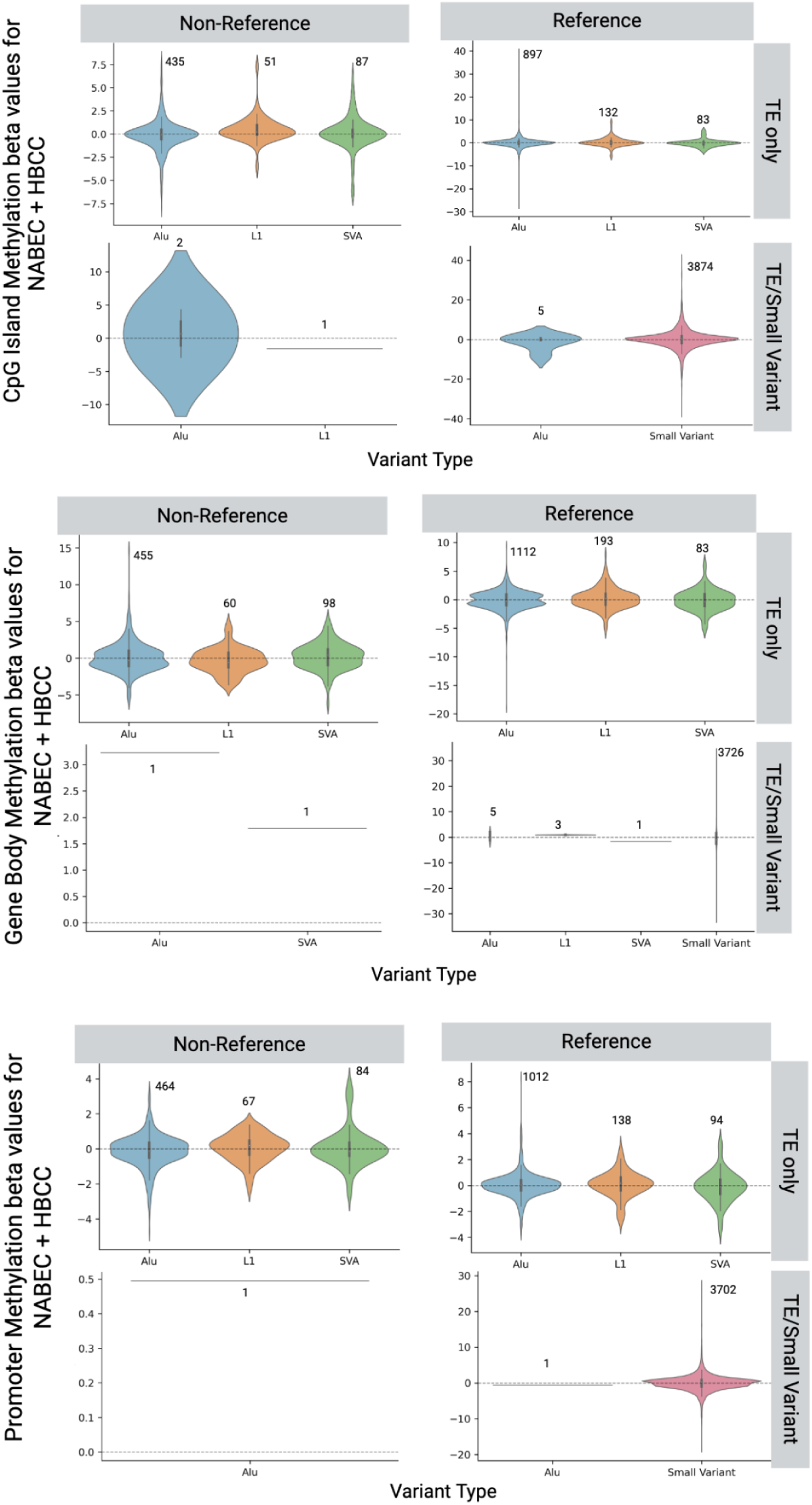
Comparison of methylation effects by variant types for CGI, Promoter, and Gene Body Regions. For each panel: Top Left: TE-only mQTL by TE type for non-reference (insertions). Top Right: TE-only mQTL by TE type for reference (deletions). Bottom left: TE-Small variant joint mQTL by TE type for non-reference (insertions). Bottom Right: TE-Small variant joint mQTL by TE type for reference (deletions).

**Supplementary Figure 5.**
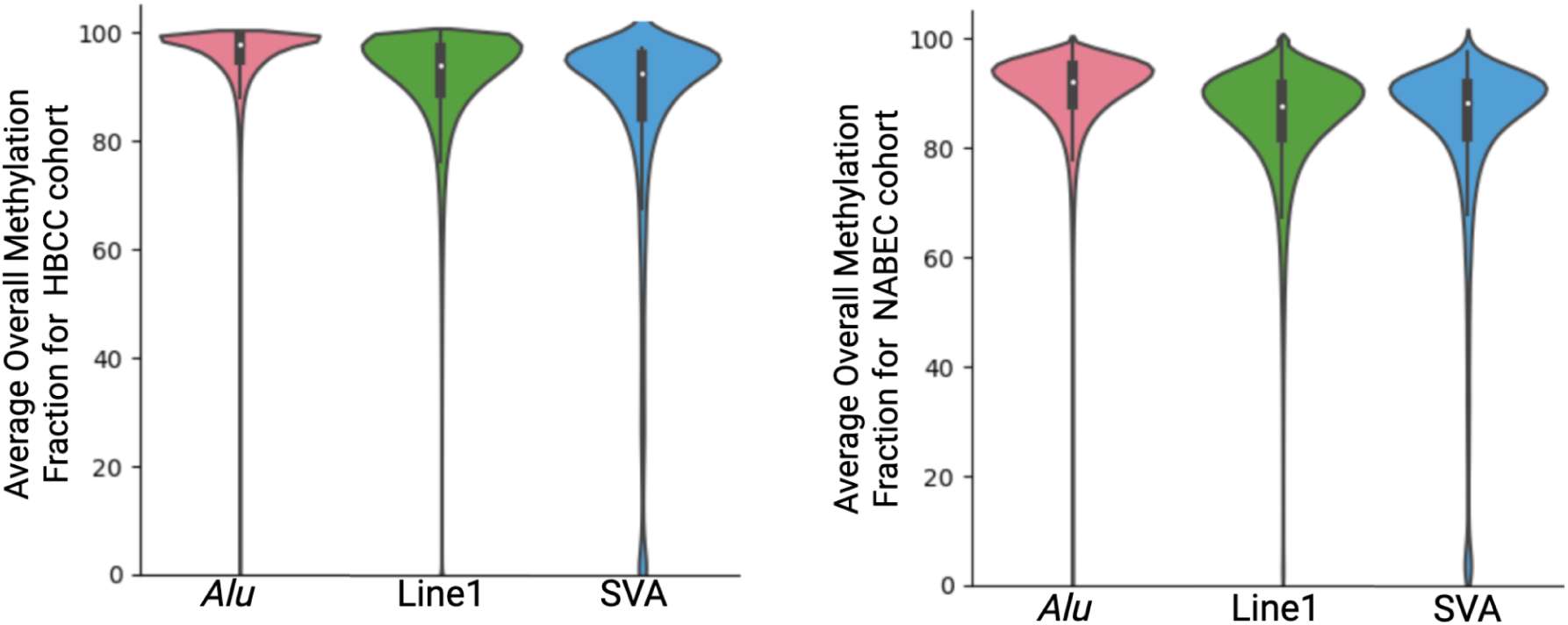
Methylation patterns of *Alu*, LINE-1, and SVA elements at CpG sites in the frontal cortex. Methylation fraction (%) for both haplotypes combined, with values >80% classified as hypermethylated and <20% as hypomethylated. Left: HBCC cohort. Right: NABEC cohort.

**Supplementary Figure 6.**
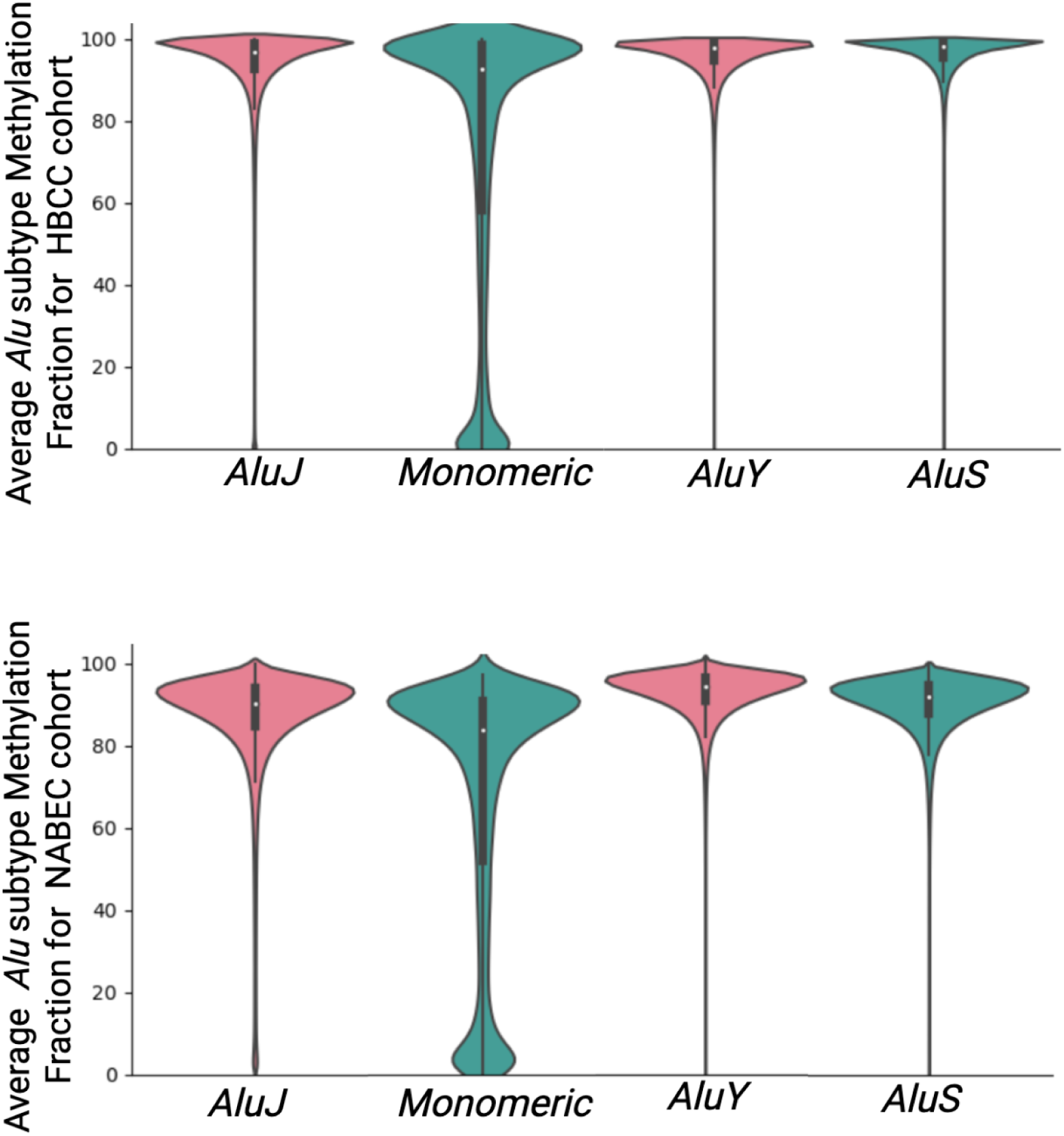
*Alu* methylation patterns by subclass at CpG sites in the frontal cortex. *Alu* subclasses include monomeric (oldest), *AluJ, AluS, and AluY* (youngest). **Top:** HBCC cohort. **Bottom:** NABEC cohort. Methylation fraction (%) for both haplotypes combined, with values >80% classified as hypermethylated and <20% as hypomethylated.

**Supplementary Figure 7.**
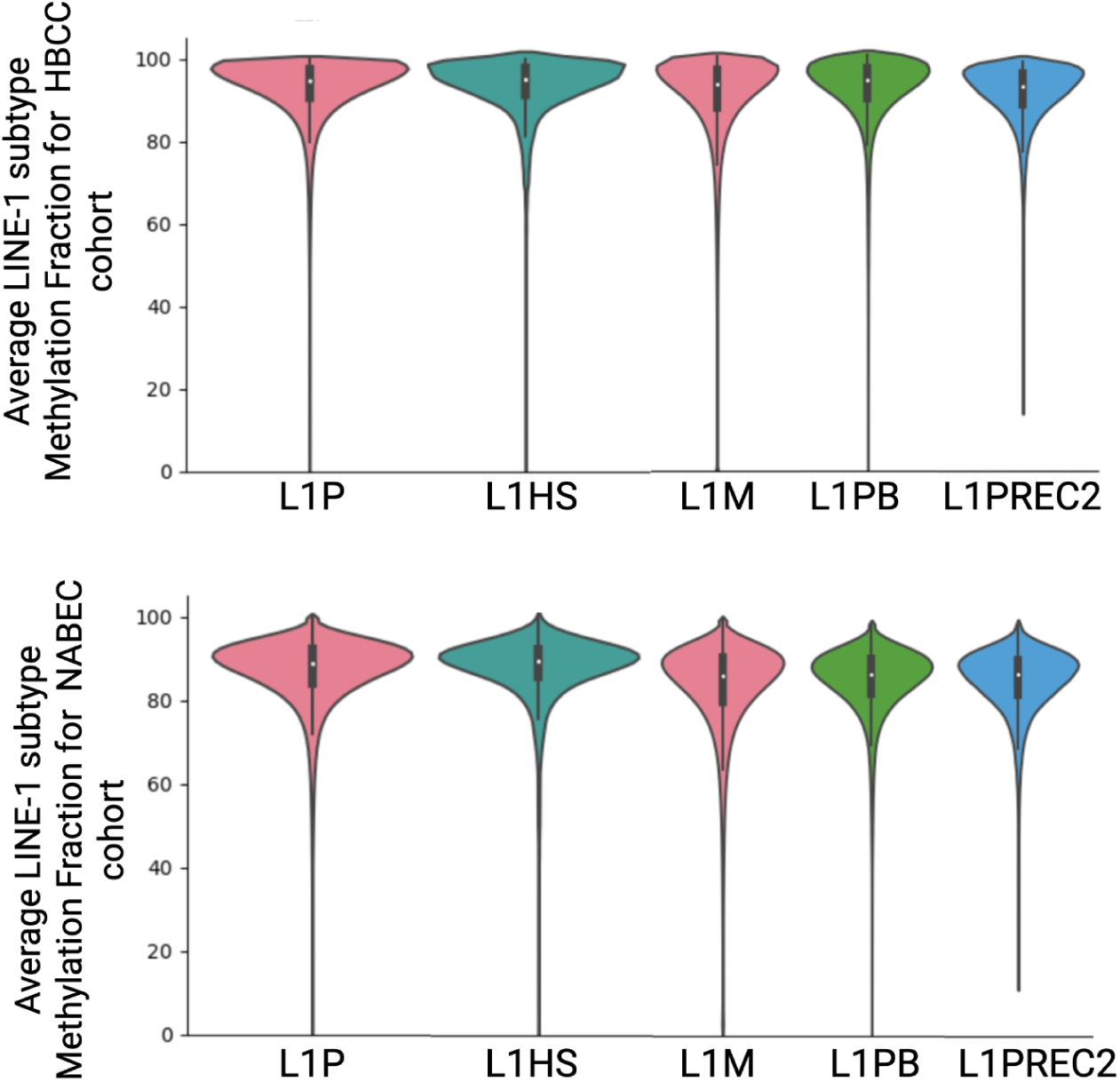
LINE-1 Methylation Patterns by CpG site in the Frontal Cortex by Subclass. Classes include LINE-1P, LINE-1M, LINE-1PB, LINE-1PREC2, and LINE-1HS (the latter being the youngest and human-specific). Top: HBCC Cohort. Bottom: NABEC cohort. Methylation fraction out of 100, in which over 80 is considered hypermethylated and under 20 is hypomethylated, for both haplotypes combined.

**Supplementary Figure 8.**
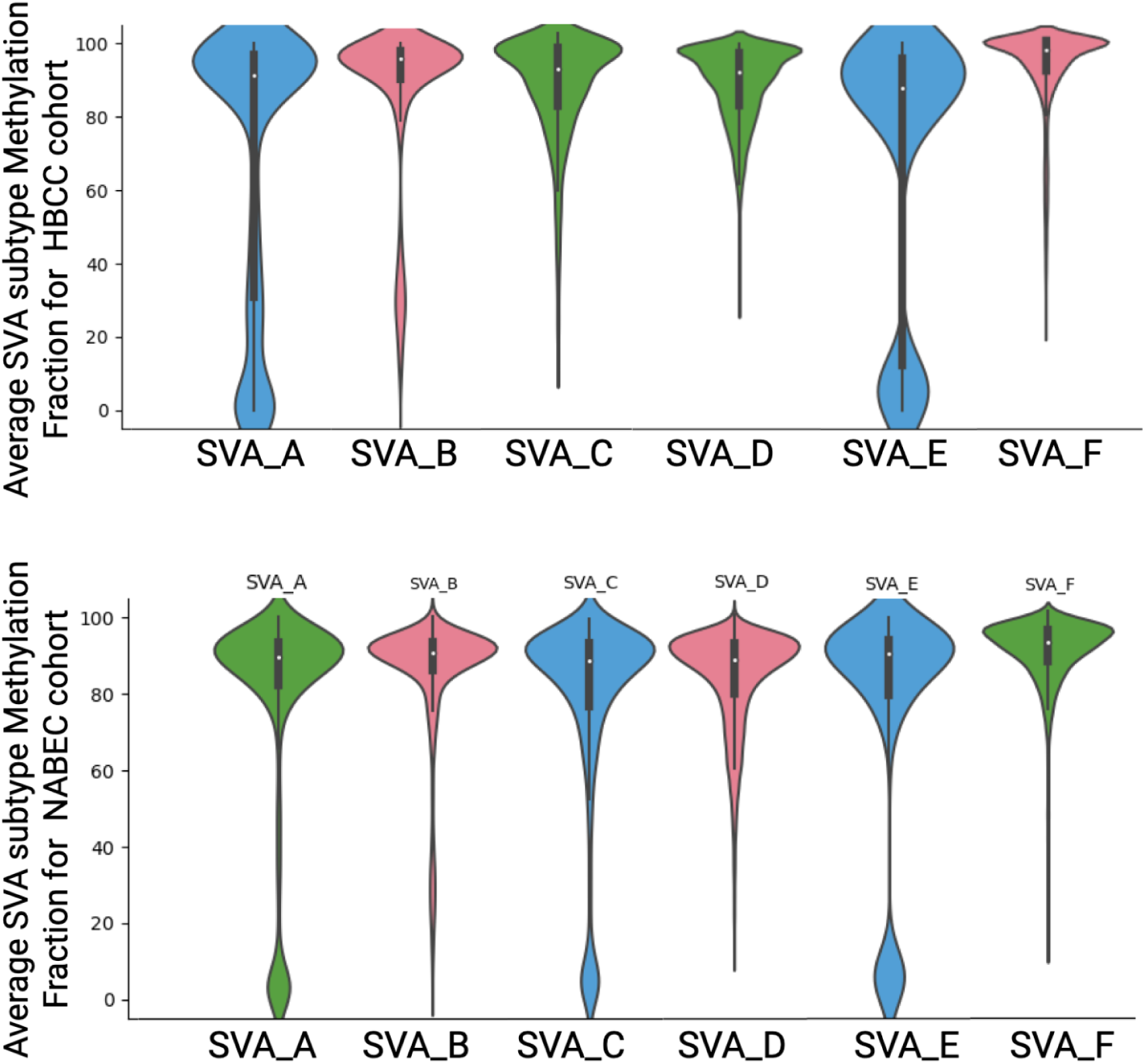
SVA methylation patterns by subclass at CpG sites in the frontal cortex. Subclasses include SVA_A–SVA_F, with SVA_F being the youngest and human-specific. Top: HBCC cohort. Bottom: NABEC cohort. Methylation fraction (%) for both haplotypes combined, with values >80% classified as hypermethylated and <20% as hypomethylated.

**Supplementary Figure 9.**
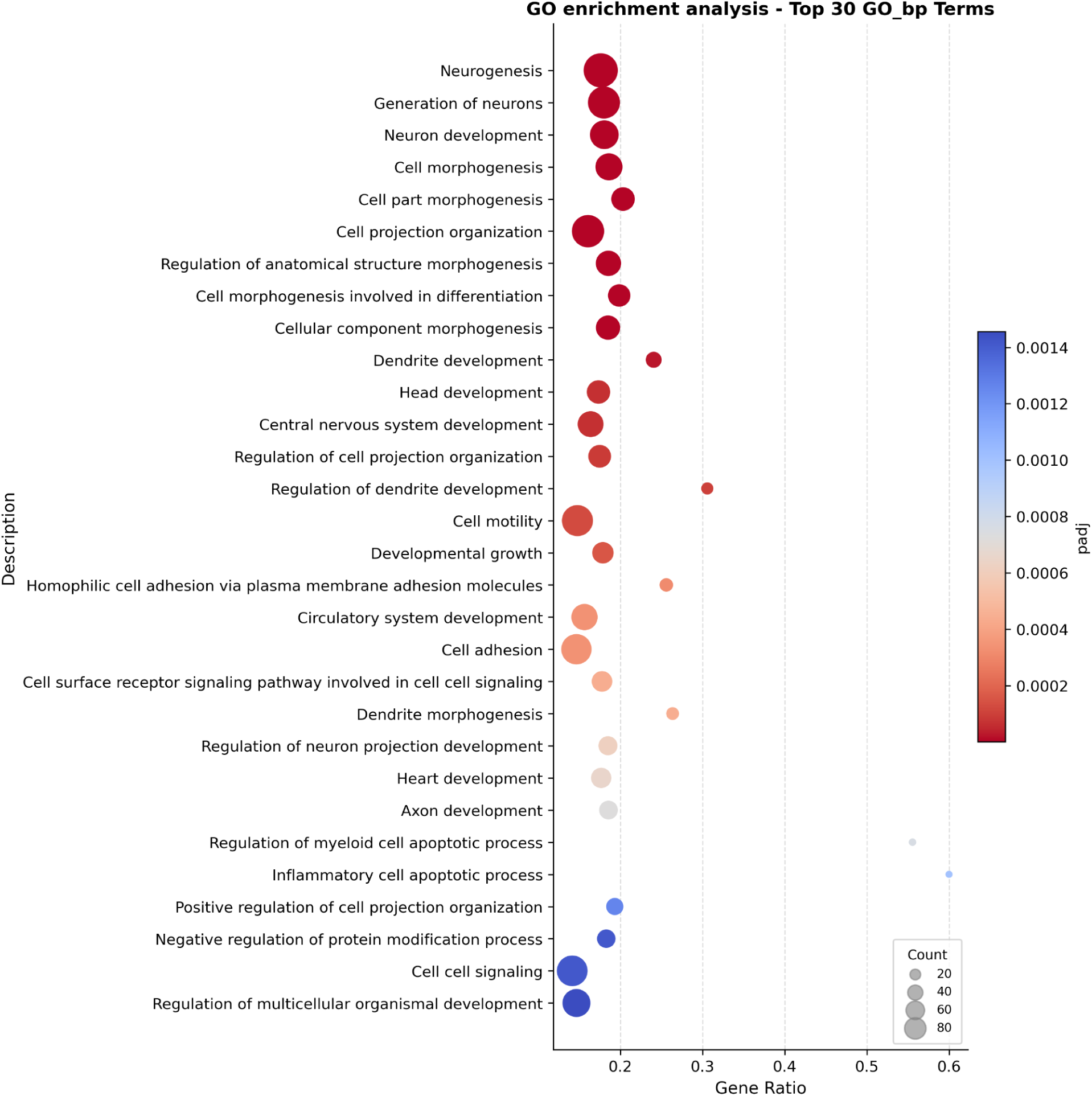
GO enrichment P-values for Biological Processes associated with NABEC-specific genes. Dot size reflects how many genes are driving that enrichment; gene ratio reflects the number of genes driving enrichment relative to the total number of genes in the GO term; and dot color reflects the p-value (significance) of the enrichment.

## Acknowledgements

We thank members of the North American Brain Expression Consortium (NABEC) for providing samples derived from brain tissue. We are grateful to the Banner Sun Health Research Institute Brain and Body Donation Program of Sun City, Arizona for the provision of human biological materials. This work was supported in part by the Intramural Research Program (IRP) of the National Cancer Institute (NCI), the National Human Genome Research Institute (NHGRI, ZIAHG200398), National Institute on Aging (NIA, ZIAAG000534, ZIAAG000538), and the Center for Alzheimer’s and Related Dementias (CARD), within the Intramural Research Program of the NIA and the National Institute of Neurological Disorders and Stroke (NINDS). The HBCC is funded by the National Institute of Mental Health (NIMH)-IRP (ZIC MH002903). M.M. was supported by NIH grant T32HG012344. We acknowledge the support of Oxford Nanopore Technologies staff in generating this data set, in particular A. Markham and J. Anderson. We also acknowledge the support of the PacBio team in generating the wet-lab protocol, in particular K. Liu, J. Burke, M. Kim & D. Kilburn. This work utilized the computational resources of the NIH STRIDES Initiative (https://cloud.nih.gov) through the Other Transaction agreement - Azure: OT2OD032100, Google Cloud Platform: OT2OD027060, Amazon Web Services: OT2OD027852. This work utilized the computational resources of the NIH HPC Biowulf cluster (https://hpc.nih.gov). Some authors’ participation in this project was part of a competitive contract awarded to DataTecnica LLC by the NIH to support open science research. M.A.N. also owns stock in Character Bio Inc. and Neuron23 Inc. This research was supported in part by the IRP of the NIH. The contributions of the NIH authors are considered Works of the United States Government. The findings and conclusions presented in this paper are those of the authors and do not necessarily reflect the views of the NIH or the U.S. Department of Health and Human Services.

## References

1. B. McClintock, The origin and behavior of mutable loci in maize. Proc. Natl. Acad. Sci. U. S. A. 36, 344–355 (1950).

2. G. Bourque, K. H. Burns, M. Gehring, V. Gorbunova, A. Seluanov, M. Hammell, M. Imbeault, Z. Izsvák, H. L. Levin, T. S. Macfarlan, D. L. Mager, C. Feschotte, Ten things you should know about transposable elements. Genome Biol. 19, 199 (2018).

3. A. Ali, K. Han, P. Liang, Role of transposable elements in gene regulation in the human genome. Life 11, 118 (2021).

4. L. Rishishwar, C. E. Tellez Villa, I. K. Jordan, Transposable element polymorphisms recapitulate human evolution. Mob. DNA 6 (2015).

5. W. S. Watkins, A. R. Rogers, C. T. Ostler, S. Wooding, M. J. Bamshad, A.-M. E. Brassington, M. L. Carroll, S. V. Nguyen, J. A. Walker, B. V. R. Prasad, P. G. Reddy, P. K. Das, M. A. Batzer, L. B. Jorde, Genetic variation among world populations: inferences from 100 Alu insertion polymorphisms. Genome Res 13, 1607–1618 (2003).

6. J. M. Deragon, P. Capy, Impact of transposable elements on the human genome. Ann. Med. 32, 264–273 (2000).

7. S. Ayarpadikannan, H.-S. Kim, The impact of transposable elements in genome evolution and genetic instability and their implications in various diseases. Genomics Inform 12, 98–104 (2014).

8. W. Zhou, G. Liang, P. L. Molloy, P. A. Jones, DNA methylation enables transposable element-driven genome expansion. Proc Natl Acad Sci U S A 117, 19359–19366 (2020).

9. E. Kejnovsky, V. Tokan, M. Lexa, Transposable elements and G-quadruplexes. Chromosome Res 23, 615–623 (2015).

10. A. Y. Du, J. D. Chobirko, X. Zhuo, C. Feschotte, T. Wang, Regulatory transposable elements in the encyclopedia of DNA elements. Nat Commun 15, 7594 (2024).

11. I. Vorechovsky, Transposable elements in disease-associated cryptic exons. Hum Genet 127, 135–154 (2010).

12. V. Fort, G. Khelifi, S. M. I. Hussein, Long non-coding RNAs and transposable elements: A functional relationship. Biochim Biophys Acta Mol Cell Res 1868, 118837 (2021).

13. S. Makino, R. Kaji, S. Ando, M. Tomizawa, K. Yasuno, S. Goto, S. Matsumoto, M. D. Tabuena, E. Maranon, M. Dantes, L. V. Lee, K. Ogasawara, I. Tooyama, H. Akatsu, M. Nishimura, G. Tamiya, Reduced neuron-specific expression of the TAF1 gene is associated with X-linked dystonia-parkinsonism. Am J Hum Genet 80, 393–406 (2007).

14. R. Douville, J. Liu, J. Rothstein, A. Nath, Identification of active loci of a human endogenous retrovirus in neurons of patients with amyotrophic lateral sclerosis. Ann Neurol 69, 141–151 (2011).

15. O. H. Tam, N. V. Rozhkov, R. Shaw, D. Kim, I. Hubbard, S. Fennessey, N. Propp, NYGC ALS Consortium, D. Fagegaltier, B. T. Harris, L. W. Ostrow, H. Phatnani, J. Ravits, J. Dubnau, M. Gale Hammell, Postmortem Cortex Samples Identify Distinct Molecular Subtypes of ALS: Retrotransposon Activation, Oxidative Stress, and Activated Glia. Cell Rep 29, 1164–1177.e5 (2019).

16. M. Prudencio, P. K. Gonzales, C. N. Cook, T. F. Gendron, L. M. Daughrity, Y. Song, M. T. W. Ebbert, M. van Blitterswijk, Y.-J. Zhang, K. Jansen-West, M. C. Baker, M. DeTure, R. Rademakers, K. B. Boylan, D. W. Dickson, L. Petrucelli, C. D. Link, Repetitive element transcripts are elevated in the brain of C9orf72 ALS/FTLD patients. Hum Mol Genet 26, 3421–3431 (2017).

17. P. Dembny, A. G. Newman, M. Singh, M. Hinz, M. Szczepek, C. Krüger, R. Adalbert, O. Dzaye, T. Trimbuch, T. Wallach, G. Kleinau, K. Derkow, B. C. Richard, C. Schipke, C. Scheidereit, H. Stachelscheid, D. Golenbock, O. Peters, M. Coleman, F. L. Heppner, P. Scheerer, V. Tarabykin, K. Ruprecht, Z. Izsvák, J. Mayer, S. Lehnardt, Human endogenous retrovirus HERV-K(HML-2) RNA causes neurodegeneration through Toll-like receptors. JCI Insight 5 (2020).

18. C. Guo, H.-H. Jeong, Y.-C. Hsieh, H.-U. Klein, D. A. Bennett, P. L. De Jager, Z. Liu, J. M. Shulman, Tau Activates Transposable Elements in Alzheimer’s Disease. Cell Rep 23, 2874–2880 (2018).

19. S. Kõks, A. L. Pfaff, L. M. Singleton, V. J. Bubb, J. P. Quinn, Non-reference genome transposable elements (TEs) have a significant impact on the progression of the Parkinson’s disease. Exp Biol Med (Maywood) 247, 1680–1690 (2022).

20. Y. Fu, G. L. Adler, P. Youssef, K. Phan, G. M. Halliday, N. Dzamko, W. S. Kim, Human Endogenous Retrovirus K in Astrocytes Is Altered in Parkinson’s Disease. Mov Disord 40, 683–692 (2025).

21. A. Fröhlich, A. L. Pfaff, V. J. Bubb, J. P. Quinn, S. Koks, Reference LINE-1 insertion polymorphisms correlate with Parkinson’s disease progression and differential transcript expression in the PPMI cohort. Sci Rep 13, 13857 (2023).

22. S. Shahid, R. K. Slotkin, The current revolution in transposable element biology enabled by long reads. Curr. Opin. Plant Biol. 54, 49–56 (2020).

23. E. J. Gardner, V. K. Lam, D. N. Harris, N. T. Chuang, E. C. Scott, W. S. Pittard, R. E. Mills, 1000 Genomes Project Consortium, S. E. Devine, The Mobile Element Locator Tool (MELT): population-scale mobile element discovery and biology. Genome Res 27, 1916–1929 (2017).

24. J. Wu, W.-P. Lee, A. Ward, J. A. Walker, M. K. Konkel, M. A. Batzer, G. T. Marth, Tangram: a comprehensive toolbox for mobile element insertion detection. BMC Genomics 15, 795 (2014).

25. J. Zhuang, J. Wang, W. Theurkauf, Z. Weng, TEMP: a computational method for analyzing transposable element polymorphism in populations. Nucleic Acids Res 42, 6826–6838 (2014).

26. P. Ramirez, W. Sun, S. K. Dehkordi, H. Zare, B. Fongang, K. F. Bieniek, B. Frost, Nanopore-based DNA long-read sequencing analysis of the aged human brain, bioRxiv (2024)p. 2024.2.0.578450.

27. K. J. Billingsley, M. Meredith, K. Daida, P. A. Jerez, S. Negi, L. Malik, R. M. Genner, A. Moller, X. Zheng, S. B. Gibson, M. Mastoras, B. Baker, C. Kouam, K. Paquette, P. Jarreau, M. B. Makarious, A. Moore, S. Hong, D. Vitale, S. Shah, J. Monlong, C. B. Pantazis, M. Asri, K. Shafin, P. Carnevali, S. Marenco, P. Auluck, A. Mandal, K. H. Miga, A. Rhie, X. Reed, J. Ding, M. R. Cookson, M. Nalls, A. Singleton, D. E. Miller, M. Chaisson, W. Timp, J. Raphael Gibbs, A. M. Phillippy, M. Kolmogorov, M. Jain, F. J. Sedlazeck, B. Paten, C. Blauwendraat, Long-read sequencing of hundreds of diverse brains provides insight into the impact of structural variation on gene expression and DNA methylation, bioRxiv (2024)p. 2024.1.16.628723.

28. C. Groza, X. Chen, T. J. Wheeler, G. Bourque, C. Goubert, A unified framework to analyze transposable element insertion polymorphisms using graph genomes. Nature Communications 15, 1–17 (2024).

29. A. E. J. van Bree, R. L. F. P. Guimarães, M. Lundberg, E. R. Blujdea, J. L. Rosenkrantz, F. T. G. White, J. Poppinga, P. Ferrer-Raventós, A.-F. E. Schneider, I. Clayton, D. Haussler, M. J. T. Reinders, H. Holstege, D. Ewing, C. Moses, F. M. J. Jacobs, A hidden layer of structural variation in transposable elements reveals potential genetic modifiers in human disease-risk loci. Genome Res 32, 656–670 (2022).

30. S. S. Arcot, Z. Wang, J. L. Weber, P. L. Deininger, M. A. Batzer, Alu repeats: a source for the genesis of primate microsatellites. Genomics 29, 136–144 (1995).

31. J. S. Beckman, J. L. Weber, Survey of human and rat microsatellites. Genomics 12, 627–631 (1992).

32. P. Balachandran, I. A. Walawalkar, J. I. Flores, J. N. Dayton, P. A. Audano, C. R. Beck, Transposable element-mediated rearrangements are prevalent in human genomes. Nature Communications 13, 1–14 (2022).

33. S. Tetter, D. Arseni, A. G. Murzin, Y. Buhidma, S. Y. Peak-Chew, H. J. Garringer, K. L. Newell, R. Vidal, T. Lashley, B. Ghetti, B. Ryskeldi-Falcon, TAF15 amyloid filaments in frontotemporal lobar degeneration. Nature 625, 345–351 (2023).

34. Genetic activation of parkin rescues TAF15-induced neurotoxicity in a Drosophila model of amyotrophic lateral sclerosis. Neurobiology of Aging 73, 68–73 (2019).

35. G. S. Wand, C. May, V. May, P. J. Whitehouse, S. I. Rapoport, B. A. Eipper, Alzheimer’s disease. Neurology 37, 1057–1057 (1987).

36. Y. Liu, T. W. Whitfield, G. W. Bell, R. Guo, A. Flamier, R. A. Young, R. Jaenisch, Exploring the complexity of MECP2 function in Rett syndrome. Nature Reviews Neuroscience 26, 379–398 (2025).

37. D.-L. Tong, R.-G. Chen, Y.-L. Lu, W.-K. Li, Y.-F. Zhang, J.-K. Lin, L.-J. He, T. Dang, S.-F. Shan, X.-H. Xu, Y. Zhang, C. Zhang, Y.-S. Du, W.-H. Zhou, X. Wang, Z. Qiu, The critical role of ASD-related gene CNTNAP3 in regulating synaptic development and social behavior in mice. Neurobiol Dis 130, 104486 (2019).

38. A. I. Soto-Beasley, R. L. Walton, R. R. Valentino, P. W. Hook, C. Labbé, M. G. Heckman, P. W. Johnson, L. A. Goff, R. J. Uitti, P. J. McLean, W. Springer, A. S. McCallion, Z. K. Wszolek, O. A. Ross, Screening non-MAPT genes of the Chr17q21 H1 haplotype in Parkinson’s disease. Parkinsonism Relat Disord 78, 138–144 (2020).

39. M. Bi, Q. Jiao, X. Du, H. Jiang, Glut9-mediated Urate Uptake Is Responsible for Its Protective Effects on Dopaminergic Neurons in Parkinson’s Disease Models. Front Mol Neurosci 11, 21 (2018).

40. N. Jansz, DNA methylation dynamics at transposable elements in mammals. Essays Biochem 63, 677–689 (2019).

41. 1000 Genomes Project Consortium, A. Auton, L. D. Brooks, R. M. Durbin, E. P. Garrison, H. M. Kang, J. O. Korbel, J. L. Marchini, S. McCarthy, G. A. McVean, G. R. Abecasis, A global reference for human genetic variation, Nature 526, 68–74 (2015).

42. A. Fröhlich, L. S. Hughes, B. Middlehurst, A. L. Pfaff, V. J. Bubb, S. Koks, J. P. Quinn, CRISPR deletion of a SINE-VNTR-(SVA_67) retrotransposon demonstrates its ability to differentially modulate gene expression at the locus. Front Neurol 14, 1273036 (2023).

43. P. A. Larsen, M. W. Lutz, K. E. Hunnicutt, M. Mihovilovic, A. M. Saunders, A. D. Yoder, A. D. Roses, The Alu neurodegeneration hypothesis: A primate-specific mechanism for neuronal transcription noise, mitochondrial dysfunction, and manifestation of neurodegenerative disease. Alzheimers Dement 13, 828–838 (2017).

44. M. Trizzino, Y. Park, M. Holsbach-Beltrame, K. Aracena, K. Mika, M. Caliskan, G. H. Perry, V. J. Lynch, C. D. Brown, Transposable elements are the primary source of novelty in primate gene regulation. Genome Res. 27, 1623–1633 (2017).

45. V. Sundaram, J. Wysocka, Transposable elements as a potent source of diverse cis-regulatory sequences in mammalian genomes. Philosophical Transactions of the Royal Society B, doi: 10.1098/rstb.2019.0347 (2020).

46. P.-É. Jacques, J. Jeyakani, G. Bourque, The Majority of Primate-Specific Regulatory Sequences Are Derived from Transposable Elements. PLOS Genetics 9, e1003504 (2013).

47. X. Zhuo, C. Tomlinson, E. A. Belter Jr, P. K. Kuntala, W. N. Saintilnord, J. Jiang, T. Lindsay, J. F. Macias-Velasco, R. S. Fulton, T. Wang, Human Pangenome Reference Consortium, Characterizing cytosine methylation of polymorphic transposable element insertions using the human pangenome resources. Genome Res 36, 1108–1124 (2026).

48. M. Xie, C. Hong, B. Zhang, R. F. Lowdon, X. Xing, D. Li, X. Zhou, H. J. Lee, C. L. Maire, K. L. Ligon, P. Gascard, M. Sigaroudinia, T. D. Tlsty, T. Kadlecek, A. Weiss, H. O’Geen, P. J. Farnham, P. A. F. Madden, J. Mungall, A. Tam, B. Kamoh, S. Cho, R. Moore, M. Hirst, M. A. Marra, J. F. Costello, T. Wang, DNA hypomethylation within specific transposable element families associates with tissue-specific enhancer landscape. Nature Genetics 45, 836–841 (2013).

49. P. Jintaridth, A. Mutirangura, Distinctive patterns of age-dependent hypomethylation in interspersed repetitive sequences. Physiol Genomics 41, 194–200 (2010).

50. B. Baker, M. Abdelhalim, K. J Billingsley, Processing human frontal cortex brain tissue for population-scale SQK-LSK114 Oxford Nanopore long-read DNA sequencing SOP v1 (2023). 10.17504/protocols.io.kxygx3zzog8j/v1.

51. H. Li, New strategies to improve minimap2 alignment accuracy. Bioinformatics 37, 4572–4574 (2021).

52. A. Catching, C. A. Weller, F. Hu, S. Bromberek, S. Abbas, K. Daida, L. Malik, B. Baker, P. K. Auluck, L. A. Screven, K. M. Andersh, K. J. Billingsley, S. Marenco, North American Brain Expression Consortium (NABEC), M. R. Cookson, K. Van Keuren-Jensen, M. A. Nalls, A. B. Singleton, C. Blauwendraat, X. Reed, Single-nucleus multiome analysis in the human prefrontal cortex identifies gene expression and cis-regulatory elements associated with aging. Cell Rep 45, 117110 (2026).

53. F. Hormozdiari, M. van de Bunt, A. V. Segrè, X. Li, J. W. J. Joo, M. Bilow, J. H. Sul, S. Sankararaman, B. Pasaniuc, E. Eskin, Colocalization of GWAS and eQTL Signals Detects Target Genes. Am J Hum Genet 99, 1245–1260 (2016).

54. A. R. Quinlan, I. M. Hall, BEDTools: a flexible suite of utilities for comparing genomic features. Bioinformatics 26, 841–842 (2010).

